# Brinkmanship and the sensory–motor limits of pursuit predation

**DOI:** 10.64898/2026.09.25.754358

**Authors:** L. Koopmans, A.M. Hein, B.T. Martin

## Abstract

Predators routinely fail to capture prey despite being larger, faster and more powerful [1–5]. Half a century ago, theory predicted that prey could evade biomechanically superior predators using brinkmanship, delaying evasive maneuvers until the last possible instant [6], but this hypothesis has never been tested in the wild. Here, using more than 2,000 hours of high-speed stereo-camera recordings of predator and prey coral reef fishes, we show that prey successfully evade 97% of attacks. However, contrary to the standard explanation that prey evade through superior maneuverability [3, 7–10], we find that prey evade by exploiting their predator’s sensory–motor delay – the interval between a stimulus reaching the predator and the onset of the predator’s motor response. Although predators partially compensate for this by forecasting prey position during interception steering, prey consistently time evasive maneuvers within a brief window just tens of milliseconds before collision, within which no viable predator trajectory can intercept them. Prey do not, however, turn at the time that maximizes miss distance, but slightly later, a bias that makes escape robust to variability in their own response timing. These findings reveal how temporal delays in perception and action place fundamental limits on the outcome of predator-prey interactions, and show that prey can exploit delays in a way that approaches the theoretical limit of escape performance.

## Main text

Across the animal kingdom, predators are generally larger, faster and more powerful than the prey they pursue [1–3]. Yet these advantages do not reliably translate into a kill. In the wild, capture success is usually below 50%, and can be far lower [4, 5]. This pattern is most extreme in water, where capture success is often below 10% even though typical predators hold a roughly fivefold speed advantage over their prey [5, 11–13]. A widely held resolution to this paradox is that prey, being smaller and therefore more maneuverable than their predators, can offset their speed disadvantage by outflanking a pursuer with a sharp turn [5, 6, 14, 15]. Despite the appeal of this hypothesis, empirical measurements of speed and maneuverability in aquatic animals imply that smaller prey are rarely maneuverable enough to offset their speed disadvantage [5, 13], leaving open the question of how prey so often escape their biomechanically superior pursuers.

A possible resolution is that the outcome of predator-prey interactions is set not by biomechanical constraints but by computational ones. To capture a fleeing prey, a predator must continuously adjust its trajectory in response to the prey’s actions using sensory–motor control [16–18]. This task is complicated by the inherent slowness of biological brains, which imposes sensory–motor delays of tens to hundreds of milliseconds between a stimulus reaching an animal’s sensory organs and the initiation of a motor response [19]. To compensate, animals are thought to employ predictive control, in which the brain forecasts both the animal’s own motion [20, 21] and the motion of its target [21–25]. Recent theory suggests that even predators using predictive control remain vulnerable during a brief window, just tens of milliseconds before collision, in which a precisely timed turn can exploit the predator’s sensory–motor delay [21]. Such brinkmanship could explain how prey so frequently escape [5], but whether wild predators use predictive control when pursuing fleeing prey, and whether prey can time [5, 26] and steer [27, 28] maneuvers precisely enough to thwart such control, remain untested. Here, we studied pursuit-evasion interactions between wild predator and prey fishes to elucidate how pursuit and evasion strategies interact in nature.

Using paired, calibrated high-speed cameras, we recorded more than 2,000 hours of full-HD stereo footage at 120 frames per second on the coral reefs of Curaçao (Fig. 1a, Methods). Cameras were directed toward persistent colonies of brown chromis, *Azurina multilineata*, a common plankton-feeding damselfish in the Caribbean that is frequently targeted by predators. From this dataset we identified 416 attacks with unambiguous outcomes, meaning the targeted prey was either captured or escaped to shelter. All were directed at brown chromis by three piscivorous fishes: bar jack (*Caranx ruber*), sand diver (*Synodus intermedius*), and grouper (*Epinephelus sp*., not identifiable to species). Thirteen attacks ended in capture (3.1%; Fig. 1a). We attempted three-dimensional reconstruction for the 108 attacks in which the prey performed an evasive maneuver (n = 99) or was captured without one (n = 9). Of these, 75 yielded three-dimensional trajectories of sufficient quality for analysis, with a median positional precision of 10 mm (see Methods for further details). Capture success in this reconstructed subset of tracks was 5.3% (4 of 75), similar to that in the full dataset.

**Fig. 1:**
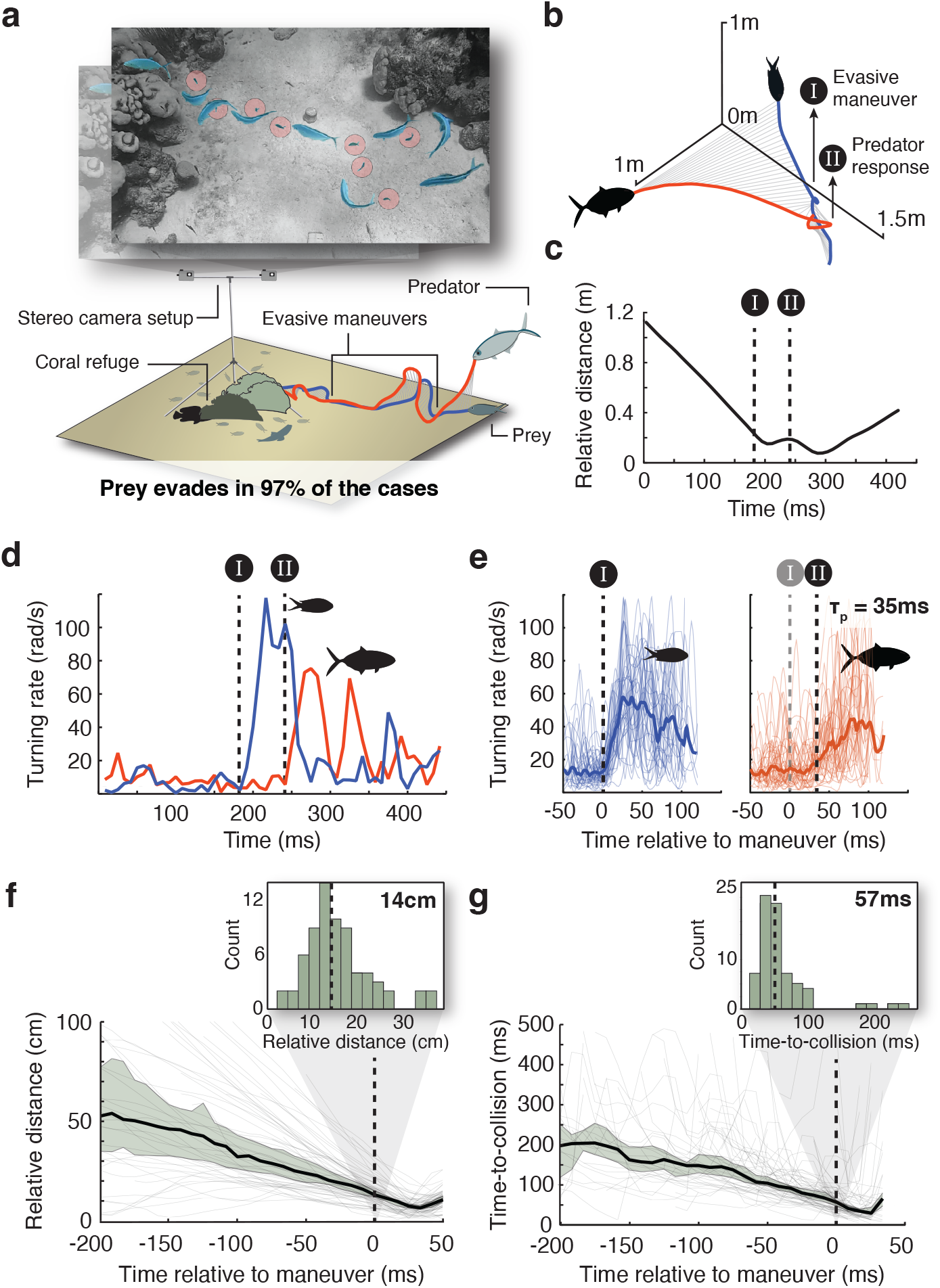
Three-dimensional reconstruction of pursuit interactions in the wild. **(a)** Top: combined still frames from a stereo recording with predator (blue) pursuing prey (red circles); Supplementary Video 1. Bottom: schematic of the stereo camera setup and 3D reconstruction of the track. **(b)** Three-dimensional reconstruction of a pursuit, with connecting lines indicating predator–prey separation over time. I marks the onset of the evasive maneuver and II the onset of the predator’s response. **(c)** Relative distance between predator and prey over time for the pursuit shown in (b); I and II as in (b). **(d)** Turning rates of predator (red) and prey (blue) for the same pursuit, with I and II as in (b). **(e)** Turning rates aligned to the onset of the prey’s evasive maneuver across reconstructed pursuits, for prey (left) and predator (right). Thin lines show individual traces; bold lines show medians. The interval between I and II is the median predator sensory–motor delay, approximately 35 ms (95% CI: 29– 48 ms, n=43, Methods). **(f)** Relative distance over time, aligned to maneuver onset, across all reconstructed pursuits. Grey lines show individual interactions; the black line shows the median; the shaded region indicates the 95% CI. Inset: distribution of predator–prey distances at maneuver onset (median = 14 cm; 95% CI: 13–16 cm; n=69; Methods). **(g)** Time-to-collision over time, aligned to maneuver onset, shown as in (f). Inset: distribution of time-to-collision at maneuver onset (median = 57 ms; 95% CI: 48–60 ms; n=69; Methods).

### Prey maneuver only when a collision is imminent

All reconstructed pursuits followed a consistent behavioral sequence. During the approach phase, predator–prey distance decreased steadily while both animals maintained low turning rates, indicating minimal changes in heading. As the predator neared the prey, the prey executed a sharp evasive maneuver that carried it away from the predator’s trajectory (Fig. 1b–d).

Predators did not respond instantaneously to these maneuvers, but only after a brief delay. We quantified this delay as the interval between the prey’s evasive turn and the onset of predator turning (Fig. 1d, time interval between I and II). Aligning predator turning-rate traces to the prey’s maneuver onset revealed a median sensory– motor delay of 35 ms (95% CI: 29–48 ms; Methods; Fig. 1e). This estimated delay is similar to previously reported sensory–motor delays in fish during visual tracking tasks (40–42 ms) [21, 29].

Prey initiated their evasive maneuvers at remarkably close range. Nearly all maneuvers were initiated within 30 cm of the predator, with a median separation of 14 cm at maneuver onset (95% CI: 13–16 cm), less than a single predator body length (mean 20.6 cm, s.d. 3.8 cm; Fig. 1f; Methods). Expressed as time-to-collision—the time until the predator would have intercepted the prey had the prey held its course—maneuvers were initiated at a median of 57 ms (95% CI: 48–60 ms), and nearly all occurred within 100 ms of predicted collision (Fig. 1g).

### Maneuvers evade predictive control

Theory suggests that a predator’s sensory–motor delay creates a brief window of opportunity for prey to escape, but the size of that window depends on the form of control the predator uses to intercept [21]. Optimal maneuver times are earlier against simple reactive control than against predictive control. Here, optimal means maximizing the miss distance, the smallest predator-prey separation during an attack. Under reactive control, the predator guides its turns directly on delayed sensory input [16]. Under predictive control, the predator compensates for its delay by estimating the sensory input it would receive in the absence of a delay, extrapolating the prey’s motion forward and recovering its own position from an internal model of its movements (Fig. 2a) [21]. To evaluate the observed maneuvers, we fitted both classes of models to the predator trajectories (Methods). Predictive control described predator trajectories better than reactive control, and both outperformed a null model in which the predator continued straight ahead (Fig. 2b,c).

**Fig. 2:**
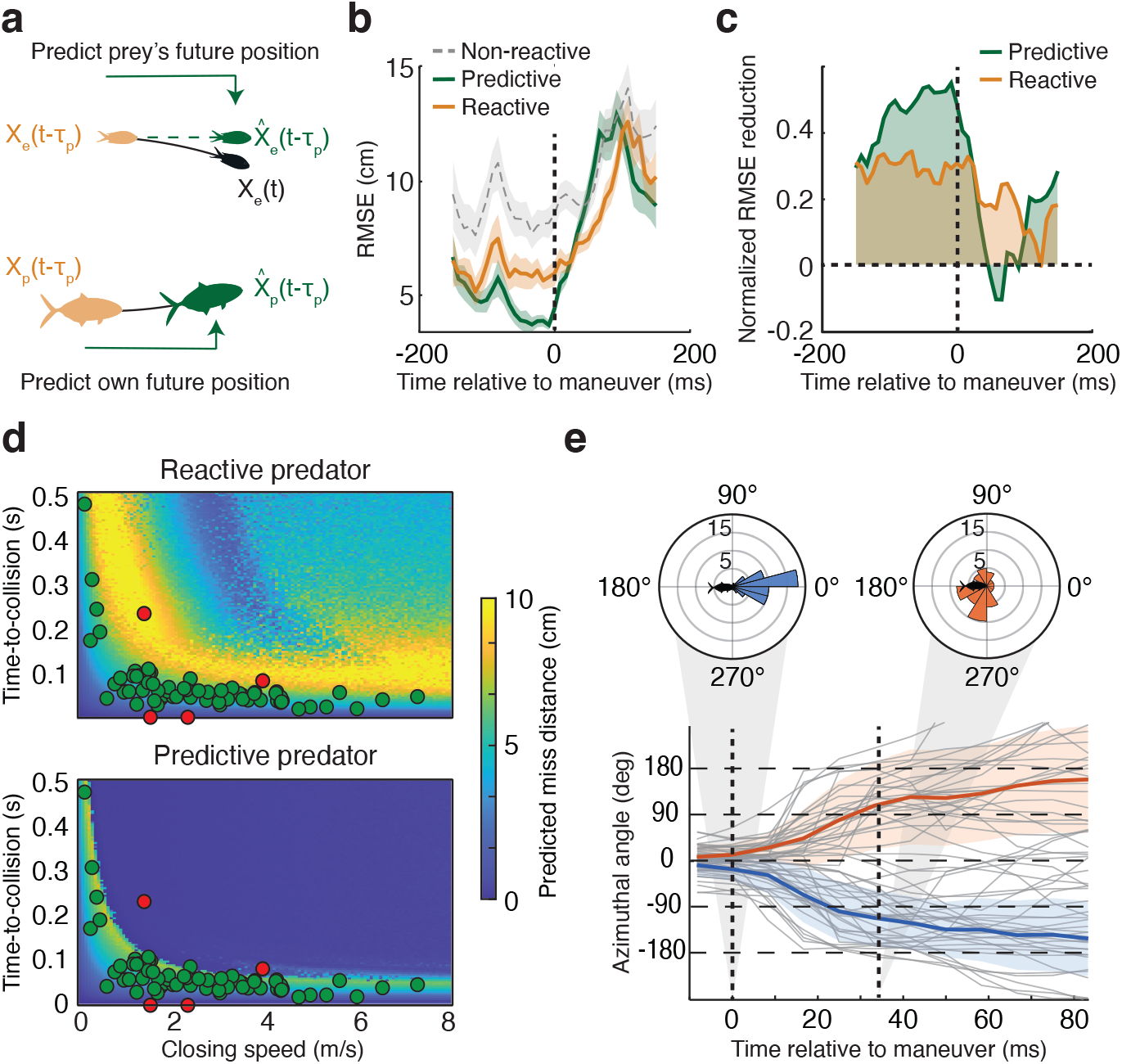
Predator and prey behavior consistent with predictive, not reactive, control. **(a)** Schematic of the two classes of interception model. A predictive predator estimates the prey’s current position 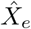 by motion extrapolation from its delayed position *X*_*e*_(*t* − *τ*_*p*_) and velocity (top), and its own position 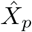 from an efference copy of its motor commands and a forward model of self-motion (bottom); a reactive predator steers on the delayed positions directly. Orange: delayed positions available at time *t*; green: predicted positions; black: the prey’s true position. **(b)** Reconstruction error (RMSE) of the predictive (green), reactive (orange) and non-reactive null (predator continues straight ahead; grey dashed) models, aligned to maneuver onset. Lines show means across attacks; shading, ±1 s.e.m. (n=71; Methods). **(c)** The same comparison expressed as the reduction in RMSE relative to the null model; shading marks the area between each curve and zero. **(d)** Predicted miss distance (color) across the closing-speed and time-to-collision plane for a reactive (top) and a predictive (bottom) predator, using the median measured prey speed, turning radii and sensory–motor delay, with predator speed set by the closing speed (*V*_*p*_ = *V*_*c*_ + *V*_*e*_). Each cell is the median over 50 escape headings drawn uniformly from 90–180°. Green circles: escapes; red circles: captures (n=71; Methods). The two captures at time-to-collision = 0 correspond to attacks where the prey did not maneuver before capture. **(e)** Azimuthal angle of the prey in the predator-centered reference frame, aligned to maneuver onset; 0° directly ahead, ±180° directly behind. Grey lines show individual maneuvers; blue and orange lines, medians ±1 s.d. for prey escaping to either side. Insets: azimuth distributions at onset and after one sensory–motor delay (dashed lines). n=54; Methods.

We then derived the relationship between maneuver timing and miss distance implied by each model (Methods). For both, the optimal maneuver time decreases with increasing closing speed, but the optimum against reactive control falls at roughly twice the time to collision of that against predictive control (Fig. 2d). Observed maneuver times fell almost entirely within the narrow window that generates substantial miss distance against predictive control, and consistently later than the optimum against reactive control. The models diverge most sharply for the maneuvers that ended in capture (Fig. 2d, red). Reactive control predicts near-maximal miss distances for these maneuvers, whereas predictive control places them outside the window that generates substantial miss distance, as observed. Several lines of evidence thus indicate that prey time their maneuvers as though facing a predator that compensates for its own delay.

Interestingly, over a brief window of roughly 100 ms following the predator’s delayed response, the fit of the predictive model (as well as that of the reactive model) collapsed to that of the null (Fig. 2c), implying that during this period, predators no longer guided their motion through visual tracking of prey. We hypothesized that this was because the prey’s maneuver moved it into a blind spot within the predator’s visual field, thereby making it impossible for the predator to continue visually-guided pursuit. To test this, we analyzed where prey were located in the predator’s visual field during the period leading up to, and following the prey’s maneuver (Methods). At maneuver onset the prey were almost directly in front of the predator (Fig. 2e). By the time the sensory–motor delay had elapsed, the prey’s turn and the predator’s own forward momentum had carried it to roughly 90° in azimuth, well outside the ∼30° frontal binocular field of teleost fishes, and it continued rearward over the following tens of milliseconds toward the caudal blind zone (Fig. 2e) [30, 31]. The maneuver may therefore disrupt the visual feedback on which continuous pursuit depends.

### No alternative predator strategy could intercept well-timed prey maneuvers

Against the best-fitting control law, prey turn at close to the optimal moment. But this leaves open whether a predator using a different guidance law, different control parameters, or a better forecast could have intercepted prey that turned at the observed times. We therefore asked a question that does not depend on the predator’s control rule: given when the prey turned, can any trajectory feasible under the predator’s sensory–motor and biomechanical constraints reach it? The delay-extended turning gambit answers exactly this question [5, 32].

In this model, both animals move at fixed speed, and neither turns more sharply than its minimum radius allows. When the prey turns, the predator holds its heading for its delay *τ*_*p*_, then turns at its maximum rate. Because the pursuer’s sharpest turn traces the boundary of the region it can reach at each instant, a prey lying outside that reachable set cannot be intercepted by any feasible trajectory (Fig. 3a; Methods). We parameterized the model with the median prey speed, turning radii and delay measured during maneuvers, with predator speed following from the closing speed on each axis (Fig. 1e; Methods). Across the observed range of closing speeds, the model predicts a window of initiation distances that no feasible predator trajectory can intercept (Fig. 3b, c). Nearly all observed successful maneuvers fall inside this region, and the captures fall outside it.

**Fig. 3:**
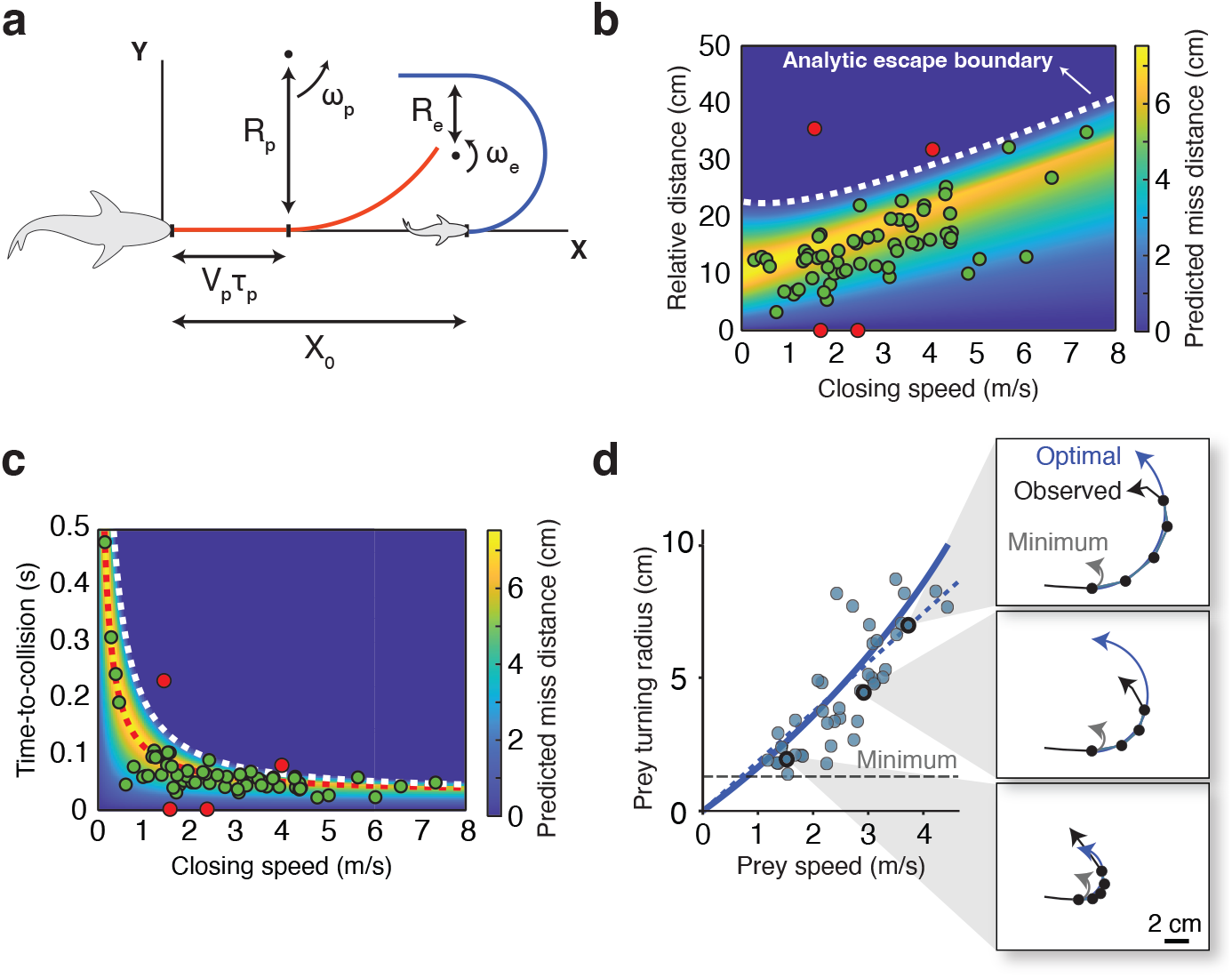
The delay-extended turning gambit predicts maneuver timing and turn radius. **(a)** Schematic of the delay-extended turning gambit. A faster predator (left) pursues a slower prey (right) head-to-tail, separated by the initiation distance *X*_**0**_. At *t* = 0 the prey turns with turning rate *ω*_*e*_ and minimum turning radius *R*_*e*_ (blue); the predator holds its heading for the duration of its sensory–motor delay, covering *V*_*p*_*τ*_*p*_, before turning with rate *ω*_*p*_ and minimum radius *R*_*p*_ (red). **(b)** Predicted miss distance (color) as a function of closing speed and predator–prey distance at maneuver onset, using the median measured prey speed, turning radii and sensory– motor delay, with predator speed set by the closing speed (*V*_*p*_ = *V*_*c*_ +*V*_*e*_). White dashed line: the analytic escape boundary; no feasible predator trajectory reaches a prey that maneuvers below it. Green circles: escapes; red circles: captures (n=66; Methods). **(c)** The same prediction in terms of closing speed and time-to-collision. White dashed line: escape boundary; red dotted line: the analytic optimum, the timing that maximizes miss distance (Supplementary Note 1). Points as in (b). The spatial escape region widens with closing speed while the temporal window contracts sharply, so faster attacks demand greater timing precision from the prey. **(d)** Prey turning radius as a function of prey speed (blue circles, n=42; Methods). The blue solid line shows the analytic optimum, the turning radius that maximizes miss distance at the median predator speed, predator turning radius and sensory–motor delay, and the blue dotted line its linear approximation. The dashed line marks the minimum turning radius (Methods). Insets show three example turns with the tracked prey positions (black points) and the optimal, observed and minimal turning arcs.

One pursuit was particularly informative about the cost of mistiming. This attack was unique in that the prey was pursued by two predators, the second trailing the first. As the leading predator closed in, the prey initiated a maneuver that fell within the escape window and carried it behind that predator (Extended Data Fig. 1; Supplementary Video 2). The same maneuver, however, was premature with respect to the trailing predator, which captured the prey.

The escape window predicted by the delay-extended turning gambit coincides closely with the one derived from the fitted predictive-control model, whereas the reactive model predicts a substantially wider window centered on earlier maneuver times (Fig. 2d). This agreement is not a coincidence: because the maneuver shifts the prey’s predicted position abruptly, the turn commanded by the predictive controller exceeds the predator’s maximum turn rate, so the trajectory it produces uses the same maximum-rate turn the turning gambit assumes (Supplementary Note 4). A predator using predictive control is therefore already at the limit set by its speed, turning radius, and sensory–motor delay. No alternative interception rule would perform better against a well-timed evasive maneuver.

While an optimally timed maneuver guarantees escape in the delay-extended turning gambit model, how sharply prey turn determines how large a miss distance they are able to generate. Maneuvers that generate small miss distances may still result in capture because a maneuver can be too small to clear the mouth of an oncoming predator. The original turning gambit [6] assumes prey turn as tightly as biomechanics allows. For aquatic animals, unlike terrestrial animals [3, 33], the minimum turn radius an animal can achieve is independent of speed [5, 10], and is approximately one quarter of body length (about 13 mm for the prey in our study [10]). While the sharpest prey turns we observed were consistent with this performance bound, most maneuvers were much wider (median turn radius 45 mm; Methods). Such wider-than-minimal turns were recently predicted to optimize miss distance in the presence of sensory–motor delays, and optimal turn radius was further predicted to increase with prey speed [5]. To test these predictions, we used the delay-extended turning-gambit model to calculate the turn radius that maximizes miss distance across the range of observed prey speeds, and compared this to the observed turn radii (Fig. 3d, Supplementary Note 1). As predicted, observed turn radius increased with prey speed. More strikingly, the observed turn radii closely tracked the optimal turn radii predicted by the model across the full range of speeds, with no fitted or free parameters. Prey thus not only time their maneuvers optimally, they also modulate their turning during maneuvers to maximize their separation from the predator at the point of closest approach.

### Predator delay, not biomechanics, limits capture

Which traits, then, determine whether a pursuit ends in capture or escape? The conventional view treats pursuit predation as a biomechanical contest in which faster or more maneuverable predators should be harder to evade [3, 7–10]. Our analysis suggests otherwise. A sensitivity analysis (Methods) revealed that neither predator speed nor turning radius can substantially reduce the miss distance produced by a well-timed prey maneuver (Fig. 4a). This holds only because prey time their maneuvers near-optimally, turning when the predator is approximately one delay away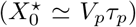— the scaling we observe across encounters (Fig. 3b, c). As a result, by the time the delay has elapsed and the predator begins to respond, it is already at the point of closest approach, and the miss distance is to first order set by the prey’s speed and the predator’s delay alone (Supplementary Note 1). Prey thus time their maneuvers so as to take the predator’s biomechanical advantage out of the equation: whatever its speed or maneuverability, the predator overshoots the prey before it can mount a response.

**Fig. 4:**
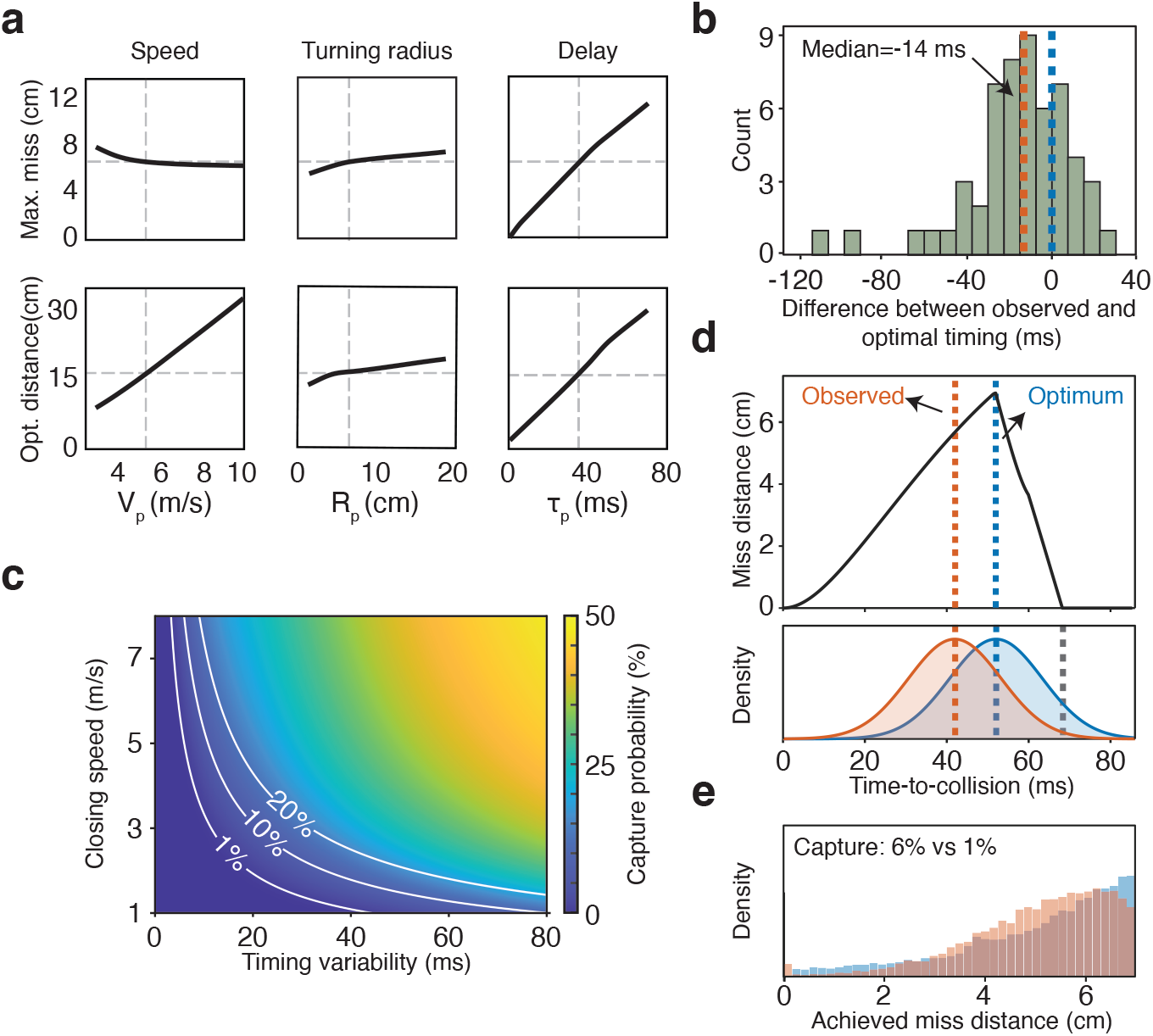
Pursuit outcomes are determined by predator sensory–motor delay and prey sensory–motor noise. **(a)** Sensitivity of escape outcomes to predator traits. Each column varies one predator trait across its observed range with the others held at their measured values, speed *V*_*p*_, turning radius *R*_*p*_, and sensory–motor delay *τ*_*p*_. For every trait value the prey chooses both when to turn and how wide to turn, so as to maximize its miss distance. Top row shows the resulting maximum miss distance, bottom the optimal initiation distance. Dashed lines mark the measured median value of each trait (vertical) and the resulting baseline prediction (horizontal). See Methods. **(b)** Histogram of difference between observed and optimal maneuver timing across the maneuvers; negative values are maneuvers initiated later than the optimum. Red dashed line: median offset (−14 ms); blue dashed line: the optimum (n=55; Methods). **(c)** Analytic capture probability across closing speed and timing variability, with 1%, 10% and 20% contours (Methods). Faster closing compresses the escape window, so the same timing variability becomes progressively more costly. **(d)** Top: predicted miss distance as a function of maneuver timing, at the median closing speed of the fastest tertile of pursuits. Bottom: realized maneuver times when timing is drawn with the measured precision of that tertile, centered on the optimum (blue) or the observed offset (14 ms later, red); grey dashed line marks the early edge of the escape window (Methods). **(e)** Achieved miss distance from the same simulation as in d; the spike at zero is capture. Aiming later reduces capture from 6% to 1% at the expense of fewer maneuvers achieving the maximum miss distance.

The only predator trait that directly controls achievable miss distance is the predator’s sensory–motor delay. As this delay approaches zero, miss distance vanishes, and a predator with instantaneous transduction from visual input to turning could intercept the prey regardless of the prey’s maneuver timing. Escape therefore depends not on outperforming the predator biomechanically, but on exploiting the time its nervous system needs to detect and respond to a turn.

### Prey hedge against their own sensory–motor noise

Observed maneuver times tracked the predicted optimum’s dependence on closing speed, but were consistently slightly later than the optimum, by a median of 14 ms (Fig. 4b) than the optimal response time. In the absence of variability in the prey’s own response timing, this bias would be suboptimal, as prey could have achieved larger miss distances by turning earlier (Fig. 4d, top). However, stochasticity in the neural circuits that control escape timing means that the exact response time varies even for identical stimulus input, with reported standard deviations of 3–20 ms depending on which sensory modality triggers the maneuver [34, 35]. A prey aiming at the optimum would therefore execute sometimes earlier and sometimes later than intended, but these deviations are not equally costly.

Early and late maneuvers fail in distinct ways: if the prey turns too late, the predator arrives before the turn and captures the prey; if it turns too early, the predator has time after its delay to adjust its heading onto a collision course with the prey. Realized miss distance decreases much more steeply for turns timed earlier than optimal than for turns that are timed later than optimal (Fig. 4d, Supplementary Note 2). A prey subject to sensory–motor noise may therefore benefit from aiming to respond slightly later than the miss-distance optimum, perhaps providing an explanation for the observed bias in prey response times relative to the optimum.

To test this, we simulated maneuvers drawn with the prey’s measured timing variability, aimed either at the deterministic miss-distance optimum or at the median observed maneuver time (Methods). Because faster closing speeds compress the window in which a maneuver can succeed, imprecision is most costly for fast-approaching predators (Fig. 4c), so we focus on the fastest third of pursuits (remaining thirds in Extended Data Fig. 2). Here, maneuvers aimed at the optimum fell outside the window more often than those aimed at the observed median offset (14 ms later, Fig. 4d, bottom). Aiming later suppressed the worst outcomes, reducing capture in these pursuits from 6% to 1%, and made intermediate miss distances more common (Fig. 4e). Together, these results suggest that the observed bias in maneuver timing is a hedge: prey sacrifice a small amount of maximum achievable miss distance for robustness against their own sensory–motor imprecision.

## Conclusions

A longstanding prediction of theoretical models of pursuit predation is that prey should engage in a brinkmanship strategy, whereby they allow their predators to approach closely before executing an evasive maneuver. Our field recordings provide the first empirical support that prey do indeed use this strategy during natural predator–prey interactions. Importantly, however, we show that the mechanism by which brinkmanship enables escape differs fundamentally from the original formulation of turning gambit theory [6]. In the original model, prey are predicted to evade by exploiting a maneuverability advantage at close distance. For this maneuverability advantage to enable escape, the prey-to-predator speed ratio must exceed the square root of the ratio of their turning radii, a condition that, based on the scaling of speed and maneuverability in the aquatic domain and typical predator–prey body size ratios, is rarely met for aquatic predator–prey pairs [5]. Instead, we find that prey time their maneuvers within a narrow window, tens of milliseconds before collision, that exploits the predator’s sensory–motor delay. A maneuver initiated in this window carries the prey behind the oncoming predator before it begins to respond, and no trajectory available to the predator can then intercept it. We furthermore find that prey steer their maneuvers so as to maximize separation within the predator’s delay: rather than turning as tightly as they can, they widen their turns at higher speeds, consistent with theoretical predictions [5].

Taken together, our results suggest that in aquatic environments, the traditional view of predator–prey interactions as a biomechanical arms race should be shifted toward a neural one. We find that even if a predator were able to increase maximum speed or turning performance, this would do little to improve its success at capturing a maneuvering prey. Instead, the ability of a predator to intercept maneuvering prey is governed primarily by the predator’s sensory–motor delay, with miss distance vanishing as this delay approaches zero. This extreme dependence of performance on sensory–motor delay may be unique to aquatic systems; due to the high density of water, aquatic prey can complete a full turn within a typical predator delay, whereas aerial and terrestrial prey rotate by only a few tens of degrees [5], so escape in those domains is likely also constrained by predator biomechanical performance [3]. Consistent with stronger selection on reaction time in water, the different fish species for which visual sensory–motor delays have been measured — trout at 42 ms [21], archer fish at 40 ms [29], and the predators studied here — fall below every other vertebrate measured [36–39]. These delays are only slightly longer than a recent estimate of the time needed for visual stimuli to be processed in the fish eye and transmitted to the brain (28 ms [40]), suggesting that there may be little room for further reducing delays through evolutionary optimization of the brain circuits that transform visual input from the retina into steering maneuvers.

Prey, in turn, are constrained by how accurately they can time their maneuvers [26]. Because the escape window is only tens of milliseconds wide, a prey that times its maneuvers optimally on average will still fall outside it as its response variability increases. The well-studied Mauthner cell (M cell) system, a neural circuit within the fish hindbrain known to produce some of the fastest behavioral responses in all vertebrates [41, 42], may provide the solution to this challenge. Not only does this circuit produce low-delay responses to sensory input (5-50 ms), it is also capable of extremely precise timing, with the standard deviation of measured sensory–motor delays in several fish species falling below 10 milliseconds [34, 43]. For prey, we expect it is more important to have precise delays than to have short delays, as a longer delay can be offset by triggering the maneuver at a correspondingly greater perceived time-to-collision [5, 13]. We find that prey further mitigate the cost of their own small sensory–motor imprecision by biasing their maneuvers later than optimal, sacrificing a small amount of miss distance for a greater tolerance to timing imprecision.

Our results provide the first empirical confirmation of brinkmanship in a natural predator-prey interaction, half a century after it was first predicted [6]. Brinkmanship succeeds because a sufficiently late maneuver makes miss distance independent of the predator’s biomechanical traits. Thus, rather than evading predators through a biomechanical trade-off between speed and maneuverability, as originally proposed, prey evade by timing their maneuvers so as to neutralize the predator’s biomechanical advantage. More broadly, our findings show that the outcome of these seemingly complex and dynamic interactions is governed by a relatively small set of biomechanical and sensory traits. This points toward the possibility of a general, predictive understanding of what drives variation in capture success across the diversity of predator–prey interactions.

## Acknowledgements

We thank Sara Neven, Lena Faber, Luna Buwalda, Doortje Nieuwenhuijzen and Eva Louwers for assistance with data collection, Mike Gil and Ronald Hassing for help developing the camera arrays, Steve Powell for acting as our local point of contact at the field site, and Caribbean Research and Management of Biodiversity (CARMABI) for storing our field equipment.

## Author contributions

L.K., A.M.H. and B.T.M. designed the study. L.K. and B.T.M. performed the field work and collected the data. L.K. and B.T.M. analyzed the data, with input from A.M.H. L.K. developed the software. L.K. wrote the initial draft of the manuscript, and L.K., A.M.H. and B.T.M. revised and approved the final version.

## Ethics

Fieldwork was conducted under the Curaçaoan Government’s Permit #2022/21467 to CARMABI. The reef sites are publicly accessible and required no further permit. The study was purely observational, with no animals captured, marked or otherwise handled, and was therefore determined not to require a license from the Animal Welfare Body of the University of Amsterdam under the Dutch Experiments on Animals Act.

## Additional information

**Supplementary Information** is available for this paper.

## Methods

### Field site and deployments

Data were collected on the leeward coast of Curaçao (12°14^′^4^′′^ N, 69°5^′^54^′′^ W) at seven sites within the bay, at depths of 3–10 m, over 38 field days in November and December 2023 (Extended Data Fig. 3). One additional attack, recorded during a trial deployment at the same site in November 2022, is included in the analyses. A deployment consisted of one stereo camera pair recording continuously. We made 347 deployments, with up to 15 pairs operating simultaneously and a mean duration of 6.1 h, totaling 0.113 petabytes of raw footage.

### Camera arrays

Each stereo pair consisted of two GoPro Hero 9 cameras in underwater housings, recording at 1920 × 1080 px and 120 fps in wide lens mode with electronic stabilization disabled to preserve a fixed intrinsic calibration. Pairs were mounted on custom-built aluminum tripods placed on the reef, with a 1 m baseline and each camera tilted inward by 5° in a cross-eyed configuration, which enlarges the volume within which three-dimensional reconstruction is possible (Supplementary Methods; Extended Data Fig. 3).

### Stereo calibration

Each stereo pair was calibrated before deployment. A checkerboard target (9 × 12 squares of 30 mm) was filmed underwater in the deployment housings at multiple orientations and distances, from under 1 m to approximately 8 m from the cameras, using a median of 50 accepted stereo pattern pairs per calibration. Intrinsic parameters (focal length, principal point, and radial and tangential distortion coefficients) were estimated for each camera separately. The extrinsic parameters, the rotation and translation between cameras, were then estimated with the intrinsics held fixed, using the Stereo Camera Calibrator app in MATLAB R2025a, which implements the method of Zhang [44]. Quality was assessed by mean reprojection error, and deployments above 1 px were excluded.

### Video synchronization

The two cameras in a stereo pair were not hardware-synchronized, so temporal alignment was performed in post-processing. The audio channels of the two streams were cross-correlated to estimate the frame offset between them, and a Python pipeline built on FFmpeg re-encoded each pair into a single side-by-side stream.

Because audio cross-correlation can leave a residual offset of up to a few frames, a further per-attack alignment was performed during reconstruction. For each annotated attack, we tested temporal offsets of ±3 frames. For each offset, we also tested both possible camera assignments, with the left and right halves of the recorded frame assigned to the left and right calibrated cameras either as recorded or swapped. We selected the combination that minimized the summed reprojection error and rectified-*y* residual. The rectified-*y* residual is the vertical offset between corresponding points after stereo rectification, which is zero for a correct geometry and so directly tests the epipolar constraint.

### Attack identification

All footage was reviewed manually. The footage was divided among three observers, amounting to about 210 hours of viewing at 10× speed. Attacks were located by two conspicuous cues, a predator crossing the frame rapidly and a collective startle response in the brown chromis colony, in which individuals dart toward cover in the coral. Footage was slowed to real time or below whenever either cue appeared. An attack was scored when a predator accelerated along a directed path toward prey, and its outcome was scored as capture, evasion, or undetermined. The “undetermined” class was assigned when the interaction left the field of view or passed behind coral before resolving. Every attack was reviewed by a second observer, who confirmed the outcome.

### Position annotation

Predator and prey positions were manually annotated using CVAT [45]. Both predator and prey were marked at the tip of the upper jaw in both camera views. Coordinates were exported as XML and parsed in MATLAB R2025a. Identical coordinates repeated in consecutive frames were treated as invalid and marked as missing. Missing positions between valid frames were filled by cubic Hermite interpolation [46].

### Stereo triangulation and track filtering

Three-dimensional trajectories were reconstructed in MATLAB R2025a from the annotated pixel coordinates, triangulating the matched left and right image points against each deployment’s stereo calibration. The resulting tracks were smoothed with a Savitzky–Golay filter [47, 48] (polynomial order 3, window 7 frames, i.e. 58 ms). A polynomial filter was chosen over a moving average because the analysis depends on derivatives (speed, turning rate and curvature) and on resolving the sharp heading change of an evasive maneuver, both of which a moving average would flatten.

### Fish body length

Body lengths were measured from stereo head- and tail-tip annotations. Predators had a mean body length of 20.6 cm (s.d. 3.8 cm; *n* = 60) and prey 5.4 cm (s.d. 1.1 cm; *n* = 41).

### Predator-centric reference frame

The predator-centered, rotation-minimizing frame used for the azimuth analysis was constructed as described in the Supplementary Methods.

### Reconstruction error and precision

Because pre-deployment calibrations were accurate to well under a pixel (0.38 px RMS, median over 56 pairs), reconstruction error reflects mainly two other sources. The first is imprecision in manual annotation, which affects every track. The second is a subset of deployments in which a camera had shifted slightly within its mount, producing a systematic vertical offset between corresponding points in the rectified image pair. Reconstruction quality was quantified by the reprojection error of the annotated prey track. Each annotated point was triangulated into three dimensions and projected back into both camera views, and its reprojection error is the pixel distance between the annotated and the projected point. These errors were summarized per track as their root-mean-square (RMS), which measures how well the two views agree on the animal’s position throughout the track. Attacks in which this error exceeded 5 px were excluded from further analysis. Reconstruction was attempted for the 108 attacks in which the prey performed an evasive maneuver or was captured without one. Of these, 82 yielded complete predator and prey tracks in both camera views, and 75 of those had a reprojection error below 5 px. All seven attacks excluded at this final step came from deployments showing the systematic offset.

Precision was quantified by Monte Carlo error propagation through the full pipeline. Smoothed three-dimensional positions were treated as ground truth, reprojected into both camera views, perturbed with isotropic Gaussian pixel noise at the measured per-track level of 1.5 px and passed back through reconstruction, giving an empirical mapping from injected noise to observed RMS reprojection error (Extended Data Fig. 4). Because error grows with distance from the cameras, as depth uncertainty in stereo triangulation scales approximately with the square of range [49, 50], precision was evaluated at each maneuver’s own distance from the camera. Median errors at maneuver onset were 1 cm in position and 0.25 m s^−1^ in speed, about 9% and 5% of the median prey and predator speeds.

### Sample sizes and inclusion criteria

Attack outcomes and capture rates by predator species are given in Extended Data Table 1. Downstream analyses restrict this pool according to the requirements of each test. The exact criteria and resulting sample size for every figure panel are given in Extended Data Table 2, and the derivation of each is described in the Supplementary Methods.

### Computation of kinematic variables

All kinematic quantities were computed from the Savitzky–Golay filtered three-dimensional trajectories (Stereo triangulation and track filtering) at a constant time step Δ*t* = 1*/*120 s.

#### Speed and turning

Velocities were estimated by three-point central difference [51, 52], with speed as their magnitude,

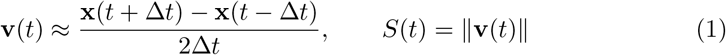

Path curvature was computed as the Menger curvature [53, 54], the reciprocal radius of the circle through three consecutive positions, which avoids differentiating position twice and is less sensitive to noise,

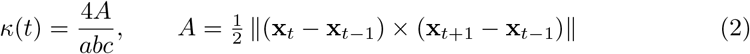

with *a* = ∥x_*t*_ − x_*t*−1_∥, *b* = ∥x_*t*+1_ − x_*t*_∥ and *c* = ∥x_*t*+1_ − x_*t*−1_∥. Turning rate follow as

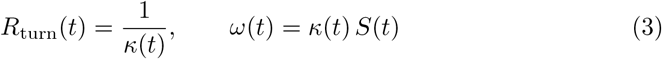

Because instantaneous curvature is sensitive to tracking noise, the turning radius of a maneuver was estimated by fitting a circle to the positions from the start of the turn, where the heading rate first exceeds 40% of its peak, until the heading had changed by 90°. These positions were projected onto their best-fit plane and fitted with an algebraic least-squares circle (Fig. 3d) [55]. Predator turns were fitted in the same way, starting one sensory–motor delay after prey onset.

### Predator–prey geometry

The relative position vector, separation and unit line-of-sight direction are

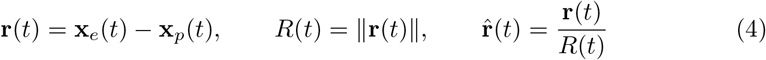

Closing speed was taken as the range rate by central difference, so that *V*_*c*_ *>* 0 indicates decreasing separation, and time-to-collision as their ratio,

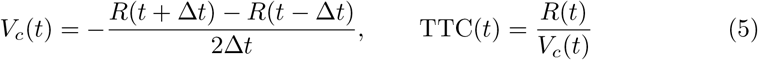

Time-to-collision was evaluated only while closing speed was positive.

The line-of-sight rate was obtained by central difference on 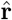,

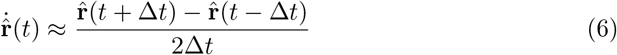

from which the line-of-sight angular velocity and its scalar rate follow,

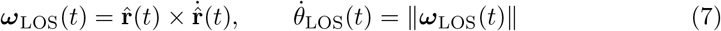

giving the scalar rate at which the line of sight rotates.

### Evasive-maneuver detection algorithm

The detector identifies the frame at which the prey initiates an evasive maneuver. The primary signal is the line-of-sight (LOS) rate 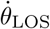, the rate at which the direction from predator to prey rotates. On a collision course the LOS does not rotate [56], so a maneuver that takes the prey off that course shows up as an increase in 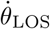. This increase builds up over several frames after the maneuver has begun, so the LOS rate itself peaks later than the onset. We therefore did not threshold the raw signal but converted it to a forward-looking rise,

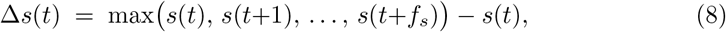

which measures how much a signal *s* increases within the next *f*_*s*_ frames and is never negative. Because it looks ahead, Δ*s* is largest at the frame where the increase starts rather than where the signal peaks, so the detector finds the start of the rise in 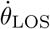 and thereby the start of the maneuver (Extended Data Fig. 5b). We used *f*_*s*_ = 4 frames (33 ms) for 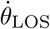.

The LOS rate can also increase when the predator turns, or when the geometry changes for other reasons. To attribute an increase to the prey, a frame was retained only if the prey’s turning rate |*ω*_*e*_| was increasing at that moment, giving the detection score

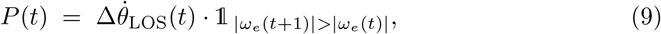

where 1_*A*_ is the indicator function, equal to 1 when condition *A* holds and 0 otherwise. Onsets were taken as local maxima of *P* (*t*) exceeding 85% of its maximum within the trial, which discards small fluctuations. Peaks less than 6 frames (50 ms) apart were merged by keeping the earlier one, so that a single continuous maneuver is not detected twice. Peaks within the first 5 frames (42 ms) of a track were discarded, because the forward-looking rise has no preceding baseline there.

### Biomechanical traits

Biomechanical traits were extracted from the reconstructed trajectories of the maneuvers that passed the reconstruction quality criterion (Field site and deployments), restricted to head-to-tail pursuits with an approach angle of at most 45 ° and a closing predator. Both evasions and captures were included, giving 64 maneuvers from 61 attacks. In the delay-extended turning gambit the outcome is set early in the encounter, while the prey turns and the predator still runs straight during its sensory– motor delay or just starts turning. The closest approach follows shortly after the delay, when the predator has turned through only a small angle (Supplementary Note 1). Speeds were therefore measured over these two phases, as the mean instantaneous speed from maneuver onset over 50 ms (6 frames) for the prey, covering most of its turn, and over 42 ms (5 frames) for the predator, covering its delay and the start of the maneuver. Turning radius was obtained from a circle fit to the first 90 ° of each turn, starting at onset for the prey and one sensory–motor delay later for the predator. Peak turning rate was the largest heading rate at the start of the turn. Because most turns are not made at an animal’s limit, we took the 90th percentile across maneuvers as its turning capacity, which stays near the top of the distribution without depending on a single noisy frame.

Median speed was 2.65 m s^−1^ for prey (95% CI 2.37–3.04 m s^−1^) and 5.15 m s^−1^ for predators (95% CI 4.94–5.47 m s^−1^). Median turning radius was 44.8 mm for prey (95% CI 33.7–50.0 mm, *n* = 42) and 62.2 mm for predators (95% CI 47.2–81.3 mm, *n* = 34). Peak turning rate was 124 rad s^−1^ for prey (95% CI 108–160 rad s^−1^) and 105 rad s^−1^ for predators (95% CI 94–116 rad s^−1^). Confidence intervals are percentile bootstrap intervals from 10,000 resamples of the maneuvers with replacement, of the median for speeds and turning radii and of the 90th percentile for peak turning rates. All computations use these values together with the sensory–motor delay.

### Sensory–motor delay estimation

The predator’s sensory–motor delay was estimated as the lag between the onset of the prey’s evasive turn and the onset of the predator’s reactive turn, using a changepoint detector applied to each animal’s absolute turning rate | *ω*| (Extended Data Fig. 6).

For each attack, *ω*(*t*) was taken from 7 frames (58 ms) before the detected maneuver onset to 10 frames (83 ms) after, and every sample past the largest value from the onset onward was discarded. Each trace therefore ends on its peak and holds only the flat baseline and the rise that the model describes. The remaining samples were upsampled fourfold by piecewise cubic Hermite interpolation [57]. That rise was modeled as

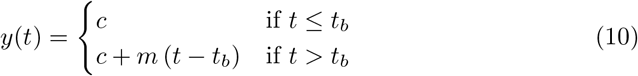

where *t*_*b*_ is the breakpoint, *c* the baseline and *m >* 0 the rising slope. For each candidate breakpoint, (*c, m*) were estimated by ordinary least squares, each segment holding at least three frames, and the breakpoint minimizing the residual sum of squares was selected. The delay was 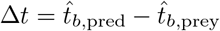.

An attack contributed only if three criteria were met. The peak turning rate after onset had to exceed 34 rad s^−1^ in both animals, twice the estimated turning-rate error propagated from the measured annotation noise (17 rad s^−1^ at the observed reprojection error of 1.5 px; Extended Data Fig. 4). The two-segment model had to improve on a flat-line null by the Bayesian information criterion. And the predator’s breakpoint could not precede the prey’s, since a response cannot come before its stimulus. The median sensory–motor delay was 35 ms (mean 39 ms; 95% CI [29, 48] ms, bootstrapped over 10,000 resamples).

### Predator guidance-law fitting

#### Delayed and predictive responses to position

Let ***p***(*t*) and ***q***(*t*) denote predator and prey position, respectively, and *τ*_*p*_ the predator’s sensory–motor delay. The reactive rule acts on delayed sensory inputs,

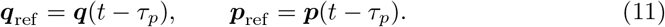

The predictive rule extrapolates prey position from the prey velocity at *t* − *τ*_*p*_ and accounts for the predator’s own displacement over [*t* − *τ*_*p*_, *t*] using an internal model of its own actions [21, 58],

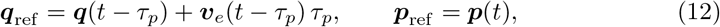

with *v*_*e*_ obtained from the reconstructed track by central differences. The two rules therefore use identical sensory information and differ only in whether that information is used as sensed or forecast. In both, the predator steers on the unit line-of-sight

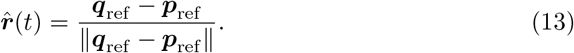

#### Guidance laws

Four steering rules were considered, each reported in pursuing animals: proportional navigation [16, 59], proportional pursuit [21], deviated pursuit [60] and parallel navigation [61]. Proportional navigation fitted best and is the family reported in the main text, in its reactive and predictive forms (Fig. 2b, c), so it is defined here. The other three are defined in the Supplementary Methods. Each rule returns a commanded turning rate *ω*_cmd_, capped at the predator’s turning capacity, *ω*_max_ = 105 rad s^−1^ (Biomechanical traits),

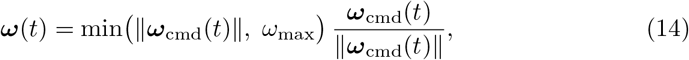

after which the heading ***û*** is advanced by rotating it about the axis ***ω****/* ∥***ω***∥ through the angle ∥***ω*** ∥Δ*t*. For proportional navigation the turning rate is proportional to the rotation rate of the line of sight,

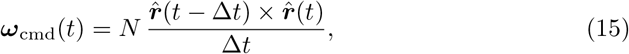

where *N* is the dimensionless navigation constant. The line-of-sight rate is formed by backward difference over one integration step, so the controller uses only information already available. This rule has been shown to be consistent with interception steering by several predatory species, including peregrine falcons [16] and predatory flies [59].

#### Sliding-window fitting of the guidance laws

Each guidance law was integrated forward from the predator’s measured position and heading at the start of the window, so a simulated track begins on the animal’s measured position. The predator was driven with its measured speed profile, so the models reproduce turning and not speed, and any later divergence reflects a mismatch between modeled and observed turning. The equations of motion were integrated by the Euler method with a step of 2.1×10^−3^ s, after upsampling the tracks fourfold from 120 to 480 Hz by piecewise cubic Hermite interpolation. *τ*_*p*_ was set to the empirically measured sensory–motor delay of 35 ms (Sensory–motor delay estimation).

Fit quality was computed in windows of 13 frames (108 ms), advanced one frame at a time within ±200 ms of maneuver onset, to follow how it changes over time. The navigation constant *N* was fitted once per track by minimizing the mean RMSE across its windows and then held fixed, so that no window is scored against a gain tuned to itself. The median fitted gain was 1.6 for predictive and 1.5 for reactive proportional navigation, so *N* = 1.5 was used for both in the simulated encounter grid below. The same procedure was applied to the other three guidance laws (Supplementary Methods).

Each window was scored by the RMSE between simulated and observed predator positions, against a non-responsive baseline of proportional navigation at *N* = 0, a predator that continues on its committed course. Windows were pooled into bins one frame wide, averaging RMSE within each track before averaging across tracks so that no track dominates a bin. Only bins with at least 15 windows were evaluated, and the band shows the standard error over tracks (Fig. 2b, c, with all eight laws in Extended Data Fig. 7).

#### Simulated encounter grid

Encounters were simulated in two dimensions over a 120 × 120 grid of closing speed *V*_*c*_ ∈ [0.1, 8] m s^−1^ and time-to-collision at maneuver onset TTC ∈ [0, 0.5] s, with a 1 ms time step (Fig. 2d). The predator starts behind the prey, heading toward it, and steers by the guidance law under test with *N* = 1.5. The prey holds course until the onset TTC is reached, then turns at its maximum rate to an escape heading. The prey speed, turning radii, and sensory–motor delay were set to their measured values (Biomechanical traits), with *V*_*p*_ = *V*_*e*_ + *V*_*c*_ and the predator’s turning rate capped at *V*_*p*_*/R*_*p*_. Each cell shows the median miss distance over 50 encounters with escape headings drawn uniformly from [90°, 180°]. Miss distance was computed as described in Computation of predicted miss distance.

#### Prey azimuth in the predator’s frame

The prey’s bearing relative to the predator was measured as the signed azimuth *ϕ* in a predator-centered, rotation-minimizing frame (Supplementary Methods). For each detected maneuver, we took the azimuth from one frame before to ten frames after onset (92 ms). Within this window, the azimuth was unwrapped by adding or sub-tracting 360° where it wrapped, so that a prey crossing directly behind the predator appears as a continuous change rather than a jump from +180° to −180°. Maneuvers were excluded if the window extended beyond the recorded track or contained missing azimuth values, leaving 54 maneuvers from 51 attacks. A prey escaping to the predator’s left increases the azimuth and one escaping to the right decreases it, so pooling both directions would average the change toward zero. Maneuvers were therefore grouped by the direction of their azimuth change and summarized per group as the median and 1 s.d. across maneuvers. The polar insets of Fig. 2e show the distribution of azimuth at onset and one sensory–motor delay later.

#### The delay-extended turning gambit

The delay-extended turning gambit is a planar geometric model of pursuit predation [5] that considers a single predator pursuing a single prey in two dimensions, an approximation supported by the near-planarity of the observed maneuvers (Supplementary Note 3). The predator is faster, but it turns on a wider circle than the prey. By turning sharply when the predator is close, the prey enters a region that the predator cannot reach without first overshooting, because its minimum turning radius does not allow it to follow the tighter turn. The predator’s sensory–motor delay extends this advantage, since it continues on its original course for the duration of the delay before it can begin to turn. The model determines, from the speeds, turning radii and delay, whether and from which initiation distances such a turn lets the prey escape. A turn started too early leaves the predator time to adjust its course, and a turn started too late ends in capture.

#### Geometry and initial conditions

We model predator and prey as point masses moving in a plane. Time *t* = 0 is the onset of the prey’s evasive maneuver. At this moment the predator is located at the origin (0, 0) and the prey at (*X*_0_, 0), so that *X*_0_ *>* 0 is the distance between predator and prey at maneuver onset, the initiation distance. Both animals initially move along the positive *x*-axis, with the predator traveling at constant speed *V*_*p*_ and the prey at constant speed *V*_*e*_ *< V*_*p*_.

Each animal *i* ∈ {*e, p*} turns at a constant angular rate *ω*_*i*_ = *V*_*i*_*/R*_*i*_, where *R*_*i*_ is its turning radius. The predator is additionally characterized by a sensory–motor delay *τ*_*p*_.

We model only the predator’s sensory–motor delay explicitly. This asymmetry reflects the geometry of pursuit: the predator can respond only after observing the prey’s evasive maneuver, whereas the prey, facing a predictable collision course, can in principle anticipate the moment of interception and time its turn accordingly.

#### Equations of motion

At *t* = 0 the prey begins a 180° turn at its minimum turning radius *R*_*e*_, which reverses its heading. The turn takes *T* = *π/ω*_*e*_, with *ω*_*e*_ = *V*_*e*_*/R*_*e*_, after which the prey swims straight. The prey’s position is then defined piecewise as

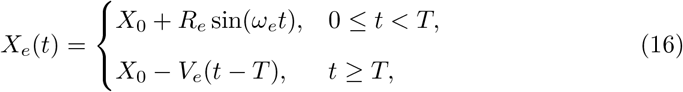

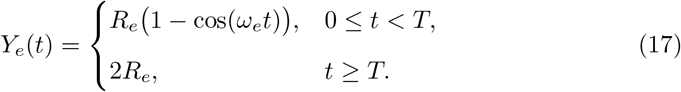

The predator trajectory incorporates the sensory–motor delay. During the delay interval 0 ≤ *t < τ*_*p*_ the predator moves straight along the *x*-direction, and for *t* ≥ *τ*_*p*_ it turns in the same direction as the prey,

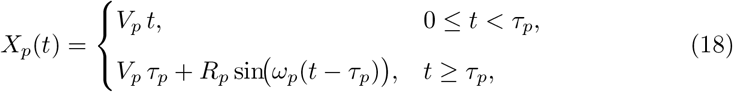

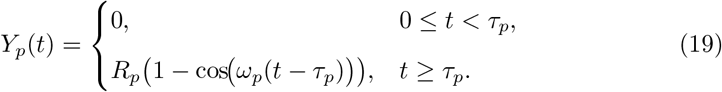

#### Computation of predicted miss distance

Writing *X*_*i*_(*t*) = (*X*_*i*_(*t*), *Y*_*i*_(*t*)) for each animal’s position, the predator–prey separation is *D*(*t*) = ∥*X*_*p*_(*t*) − *X*_*e*_(*t*)∥. Trajectories were evaluated on 5,000 points spanning *t* ∈ [0, *τ*_*p*_ + *π/ω*_*p*_], the end of the predator’s half-turn.

We located the first instant *t*_*c*_ at which the predator reaches the prey’s *x*-position, *X*_*p*_(*t*_*c*_) = *X*_*e*_(*t*_*c*_). If the prey’s lateral position then exceeds the predator’s, *Y*_*e*_(*t*_*c*_) *> Y*_*p*_(*t*_*c*_), the prey escapes [5, 6], and its miss distance *D*_min_(*X*_0_) is the smallest separation as the predator passes. If not, or if the predator does not reach the prey’s *x*-position within its half-turn, the predator can reach the prey and the miss distance is set to zero.

#### Delay-extended turning gambit predictions against observed maneuvers

The delay-extended turning gambit depends on six quantities, too many to show jointly. For Fig. 3b, c we therefore fixed prey speed, both turning radii and the delay at their measured values (Biomechanical traits), with *V*_*p*_ = *V*_*e*_ + *V*_*c*_, and varied closing speed and initiation distance. To test whether this simplification holds for individual encounters, we also computed the escape boundary of each maneuver from the prey and predator speeds measured at its onset. Both captures were initiated outside their predicted escape window and evasions within it (Extended Data Fig. 8). For Fig. 3d, each prey turning radius (Computation of kinematic variables) was plotted against the prey’s mean speed over the same circle segment, together with the radius that maximizes miss distance at the measured predator speed, turning radius and delay, and the prey’s tightest achievable radius, 13.2 mm, from the scaling reported by Domenici [10]. The escape boundary, the optimal initiation distance and the optimal turning radius are derived in Supplementary Note 1.

#### Sensitivity of escape performance to biomechanical traits

Each predator trait was varied in turn about its measured value with the others held fixed, speed over 0.5–2 ×, turning radius over 0.2–3× and sensory–motor delay over 0.01–2×, in 120 steps each. For every trait value the prey was allowed to choose both its initiation distance and its turning radius so as to maximize its miss distance, computed from the delay-extended turning gambit (The delay-extended turning gambit). The initiation distance was searched over *X*_0_ ∈ [0.001, 1.0] m and the turning radius from the tightest radius the prey can achieve, 13.2 mm, up to 150 mm, so a prey may turn wider than its minimum when that increases its miss distance. Fig. 4a shows the resulting maximum miss distance and the initiation distance at which it occurs.

#### Timing-precision analysis

We asked whether prey time their maneuvers at the optimum predicted by the delay-extended turning gambit, and whether their timing is precise enough to stay within the escape window. The analysis used the maneuvers that were not captures, had an approach angle of at most 45 ° and a closing speed of at least 0.6 m s^−1^, giving 58 maneuvers. The closing-speed floor prevents small values of *V*_*c*_ from inflating timing errors, which are obtained by dividing by *V*_*c*_.

##### Observed and optimal timing

For each maneuver, the difference between observed and optimal timing is 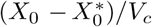, where *X*_0_ is the observed initiation distance and 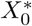 the optimal initiation distance (Supplementary Note 1) computed from that maneuver’s prey and predator speeds, with turning radii and delay at their measured values (Biomechanical traits; Fig. 4b).

##### Timing precision

Because the escape window narrows substantially with closing speed, we split the maneuvers by closing speed into three equal-sized groups. The observed maneuver timing was described by a least-squares fit of initiation distance to closing speed,

*X*_0_ = *a* + *bV*_*c*_, over all 58 filtered maneuvers. The fit was made in distance because the scatter in initiation distance does not change with closing speed. Each residual was converted to a timing error by dividing by that maneuver’s *V*_*c*_, and the timing precision *σ* of a group was taken as the standard deviation of its timing errors, with a bootstrap 95% CI from 10^4^ resamples (Fig. 4d, with all three groups in Extended Data Fig. 2).

##### Capture probability

A maneuver fails if it is made too early, beyond the escape boundary 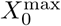, or too late, before the prey can turn. For a timing distribution centered at initiation distance *X*_*c*_, these define two margins in time, 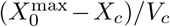 and *X*_*c*_*/V*_*c*_, evaluated at the group’s median closing and prey speeds. With Gaussian timing errors, the capture probability is the probability of falling beyond either margin (Supplementary Note 2). For each group this was evaluated at its measured *σ*, centered on the observed maneuver timing (Fig. 4d, e; Extended Data Fig. 2). Fig. 4c shows the capture probability over a 400 × 400 grid of *V*_*c*_ ∈ [1, 8] m s^−1^ and *σ* ∈ [0, 80] ms, centered on the optimum.

#### Miss-distance distributions

To see how timing errors affect the miss distance and not only the risk of capture, we drew 20,000 maneuver times from a Gaussian at each group’s measured *σ* and median closing speed. The draws were centered once on the optimal timing and once on the observed maneuver timing, so the two differ only in where the prey times its maneuver. For each drawn maneuver time, the miss distance was computed from the delay-extended turning gambit (The delay-extended turning gambit), with a capture counted as a miss distance of zero (Fig. 4d, e, with all three groups in Extended Data Fig. 2).

## Data availability

The reconstructed three-dimensional predator–prey trajectories, detected maneuver times and derived kinematic variables that support the findings of this study are available at Zenodo (https://doi.org/10.5281/zenodo.22947421)

## Code availability

All analysis code is available at Zenodo (https://doi.org/10.5281/zenodo.22947421)

## Extended Data

**Fig. 1: Extended Data Fig. 1.**
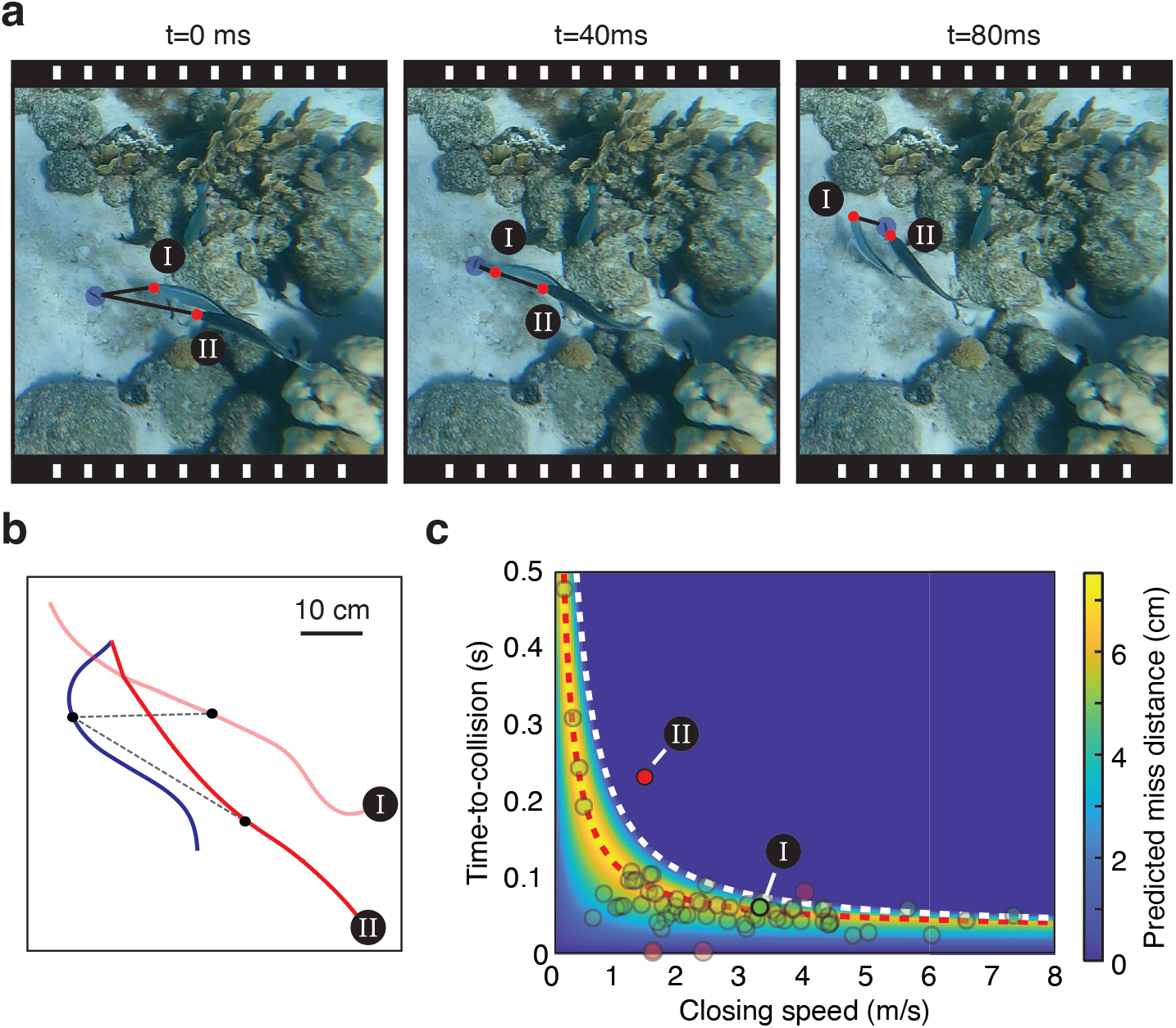
One maneuver against two pursuing predators. **(a)** Video frames Supplementary Video 2 of a pursuit in which two bar jacks chased the same prey, the second trailing the first. Blue marker: prey. Red markers: the two predators, I leading and II trailing. The prey’s turn carried it behind predator I, which it evaded, while predator II captured it. **(b)** Reconstructed tracks of the same pursuit seen from above, in top-down view. Blue: prey. Light red: predator I. Red: predator II. Black points mark the positions at maneuver, and dashed lines the separation to each predator at that moment, 19.6 cm to predator I and 35.6 cm to predator II. The tracks end where predator II reaches the prey, 67 ms after onset. **(c)** The same maneuver placed twice in the turning-gambit plane of Fig. 3c, once for each predator, since each was at its own distance and closing speed at that instant. Color: predicted miss distance at the measured prey speed, turning radii and sensory– motor delay, with predator speed set by the closing speed. White dashed line: the escape boundary. Red dotted line: the timing that maximizes miss distance. Transparent circles: the other observed maneuvers. Relative to predator I the turn came at a time to collision of 58 ms, 3 ms from that encounter’s optimum of 55 ms and well inside its escape window, with a predicted miss distance of 8.6 cm. Relative to predator II the same turn came at 229 ms, 119 ms earlier than that encounter’s optimum of 111 ms and beyond its escape boundary of 196 ms, where the model predicts interception. The maneuver timed for one pursuer is therefore premature for a second one that is farther away and closing more slowly (Methods).

**Fig. 2: Extended Data Fig. 2.**
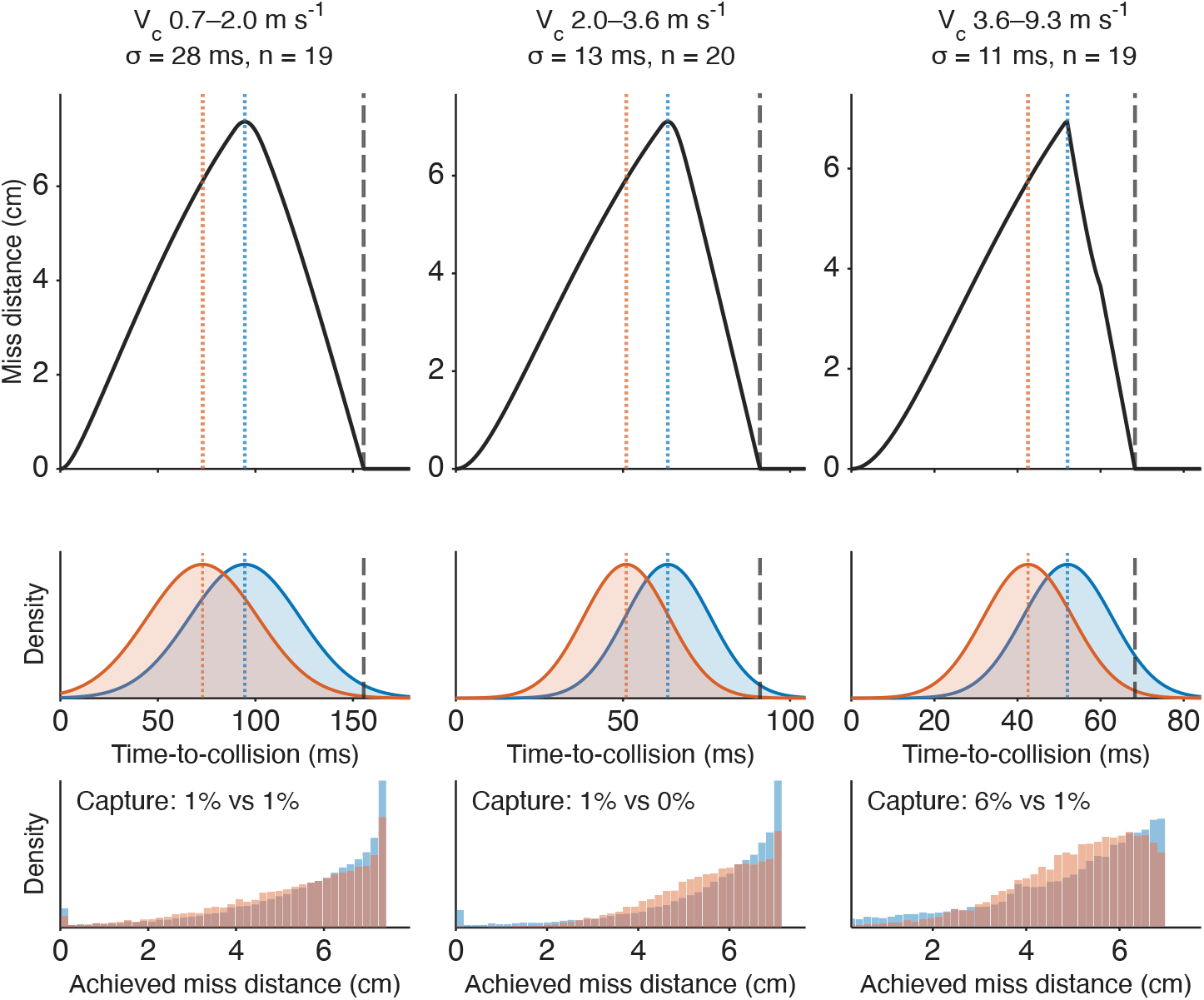
Miss distance under timing noise for all closing-speed groups. Escapes were split by closing speed into three equal-sized groups (columns, slowest to fastest), each labeled with its closing-speed range, timing precision *σ* and number of maneuvers. The right-hand column is the group shown in Fig. 4d, e. Top, predicted miss distance against the time-to-collision at which the prey turns, evaluated at the group’s median closing speed. Dotted lines mark the optimal (blue) and observed (red) maneuver timing, and the dashed gray line the escape boundary, beyond which the prey turns too early to escape. Middle, maneuver times drawn from a Gaussian with the group’s *σ*, centered on the optimal or the observed timing. Bottom, the resulting miss distances, with captures counted as zero. Capture probability was lower for observed than for optimal timing in every group (0.7 vs 1.5%, 0.1 vs 1.4% and 0.7 vs 6.5%, slowest to fastest).

**Fig. 3: Extended Data Fig. 3.**
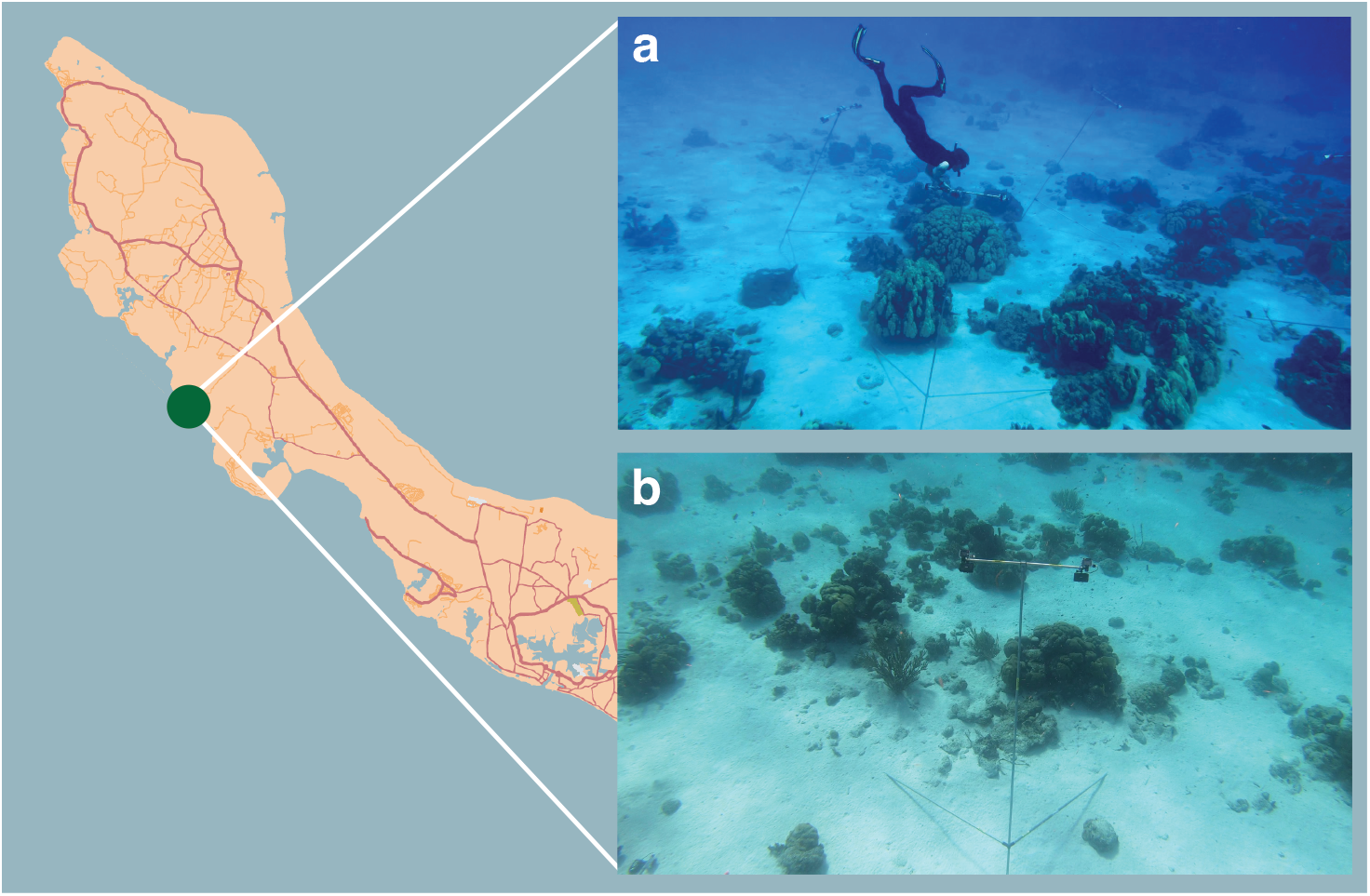
Field site and camera arrays. All recordings were made at a single reef site on the leeward (west) coast of Curaçao, southern Caribbean (green circle on the map). **(a)** Camera arrays deployed on the reef, with four stereo pairs visible. Up to 15 stereo camera pairs were deployed on a single day. Each pair was positioned to view persistent colonies of brown chromis (*Azurina multilineata*). **(b)** A single stereo pair mounted on a custom-built aluminum tripod, consisting of two 2 m horizontal rods resting on the seafloor and a slightly tilted 3 m vertical rod that placed the cameras approximately 2.5 m above the substrate. Each pair comprised two GoPro Hero 9 cameras separated by a 1 m baseline and tilted inward by 5° in a cross-eyed configuration, which maximizes binocular overlap over the 2–5 m range at which interactions were typically observed. All footage was recorded at 1920 × 1080 px and 120 frames s^−**1**^.

**Fig. 4: Extended Data Fig. 4.**
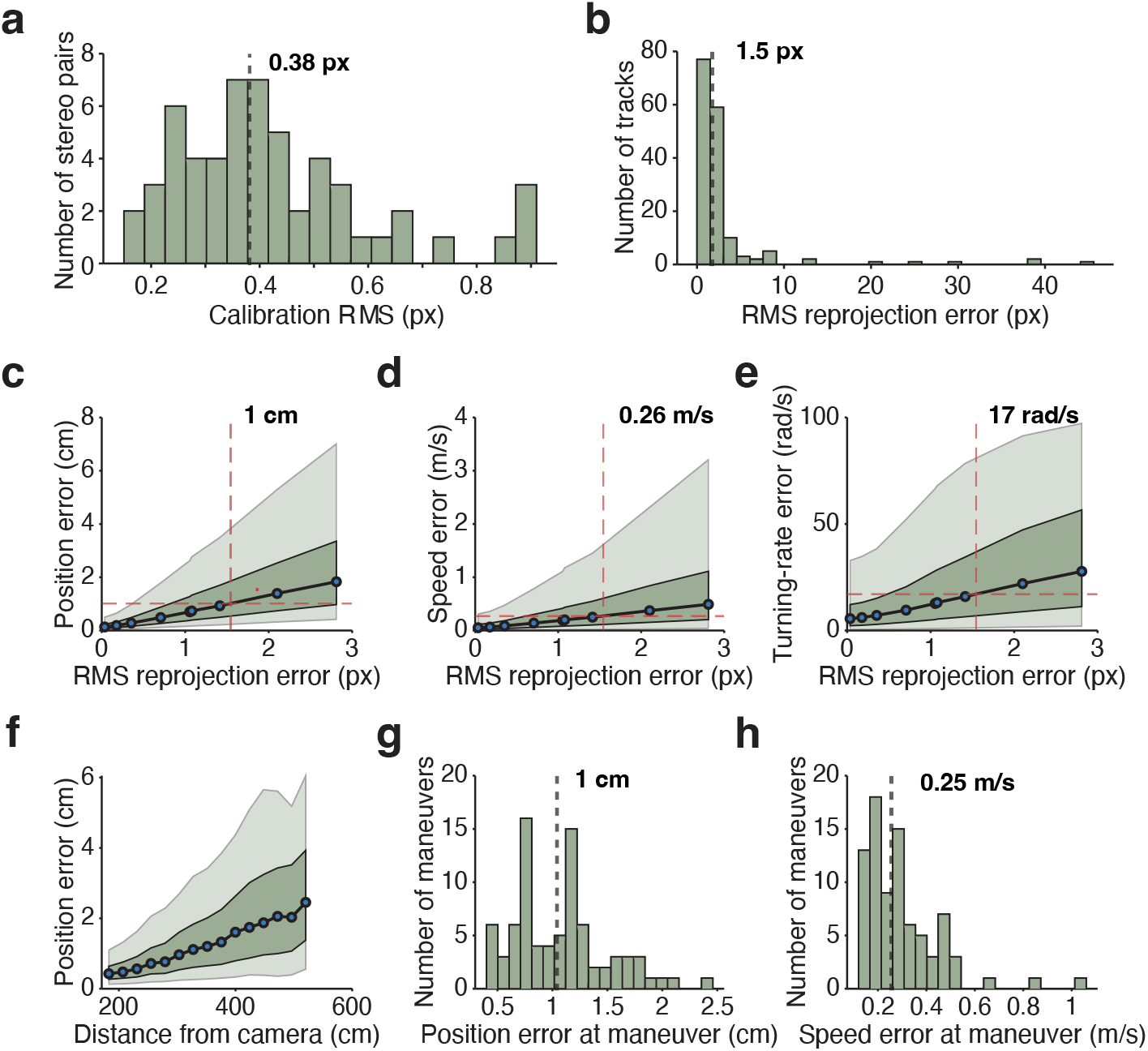
Monte Carlo error propagation through the stereo reconstruction pipeline. Smoothed three-dimensional tracks were treated as ground truth, reprojected into both camera views, perturbed with isotropic Gaussian pixel noise (*σ* = 0.05–4 px per coordinate) and passed back through the full reconstruction pipeline; 5 replicates were run at each noise level and pooled across all tracks. **(a)** Calibration RMS reprojection error across 56 stereo camera pairs, reflecting the residual of the stereo camera model alone (median 0.38 px). **(b)** Per-track RMS reprojection error, which additionally includes annotation imprecision, pooled over predator and prey tracks (median 1.51 px); tracks exceeding 5 px were excluded from the dataset. **(c–e)** Position, speed and turning-rate error as a function of the RMS reprojection error produced by the simulated annotation noise. Dashed lines mark the empirical median (1.5 px); lines show medians, with dark and light bands spanning the interquartile and 5th–95th percentile ranges. **(f)** Positional error binned by distance from the camera at the empirical noise level; the depth precision of stereo triangulation degrades with range, so the same pixel error yields a larger three-dimensional error further away. **(g**,**h)** Position and speed error at the moment of maneuver initiation, inferred from each maneuver’s own distance from the camera (medians 1 cm and 0.25 m s^−**1**^).

**Fig. 5: Extended Data Fig. 5.**
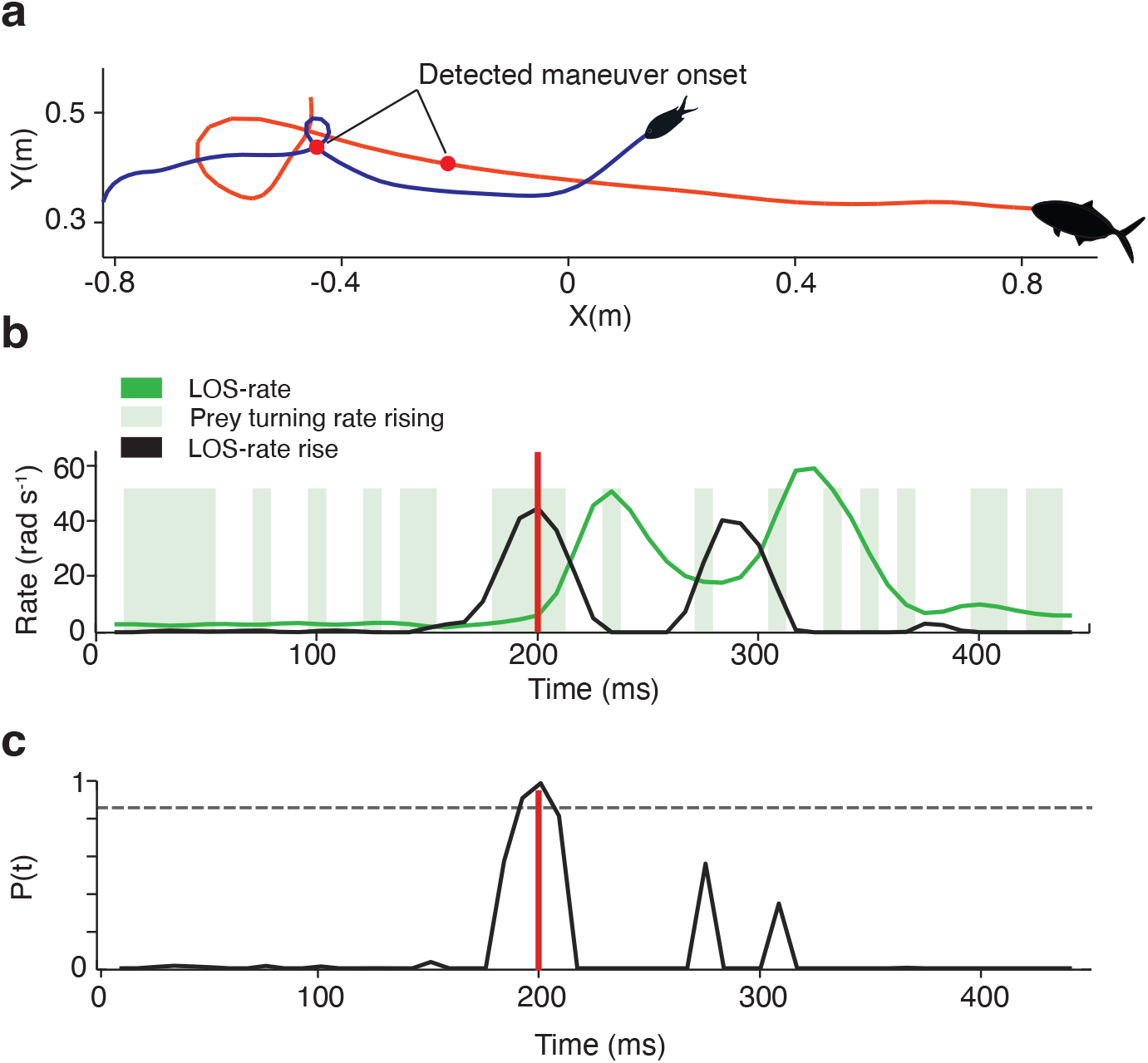
Automated detection of evasive-maneuver onset. The line-of-sight (LOS) rate 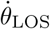 was converted to a forward-looking rise, the amount it climbs over the next 4 frames (33 ms), which is largest where the rate starts to climb rather than where it peaks. A frame was retained only while the prey’s turn rate was rising, |*ω*_*e*_(*t*+1) | *>* |*ω*_*e*_(*t*) |, and onsets were taken as local maxima of the resulting detection score exceeding 85% of its trial maximum (Methods). Data are from one example pursuit. **(a)** Top-down view of the reconstructed predator (red) and prey (blue) trajectories; filled points mark the positions of both animals at the detected maneuver onset. **(b)** LOS rate (green) and its forward-looking rise (black). Green bands mark frames in which the prey’s turn rate is rising; the red line marks the detected onset, which falls where the LOS rate starts to climb. **(c)** Detection score, the LOS rise is retained only where the prey’s turn rate is rising. The dashed line marks the 85% threshold; only the peak at ∼200 ms exceeds it, while the other candidate peaks are discarded.

**Fig. 6: Extended Data Fig. 6.**
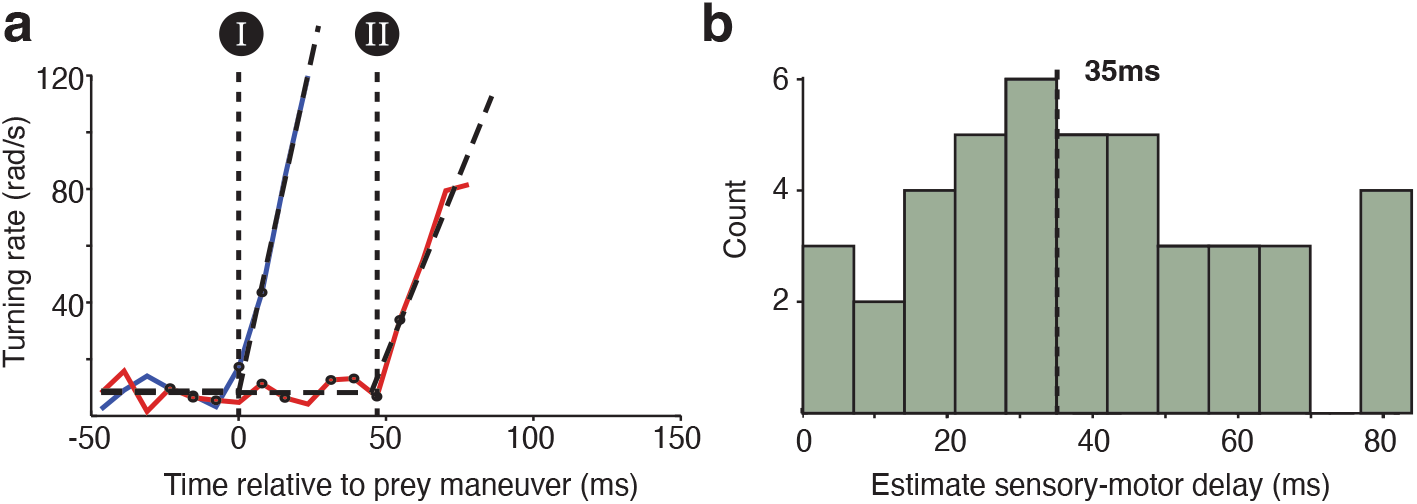
Estimation of the predator’s sensory–motor delay. The delay was estimated as the lag between the onset of the prey’s evasive turn and the onset of the predator’s reactive turn, obtained by fitting a two-segment piecewise-linear model (flat baseline followed by a linear rise) to each animal’s absolute turning rate and taking the difference between the fitted breakpoints. Attacks were retained only where the post-onset peak turning rate exceeded 34 rad s^−**1**^ in both animals, twice the turning-rate error at the measured annotation noise (Extended Data Fig. 4), the two-segment model improved on a flat-line null by the Bayesian information criterion with a positive fitted slope, and the predator’s breakpoint did not precede the prey’s. **(a)** Example fit for a single attack, showing the absolute turning rate of prey (blue) and predator (red) with the fitted models as dashed curves. Dashed vertical lines mark the prey breakpoint (I) and the predator breakpoint (II), and the interval between them is the estimated delay. **(b)** Distribution of estimated delays across the 43 attacks that passed every criterion, with a median of 35 ms (dashed line) and a 95% confidence interval of 29–48 ms from 10,000 bootstrap resamples.

**Fig. 7: Extended Data Fig. 7.**
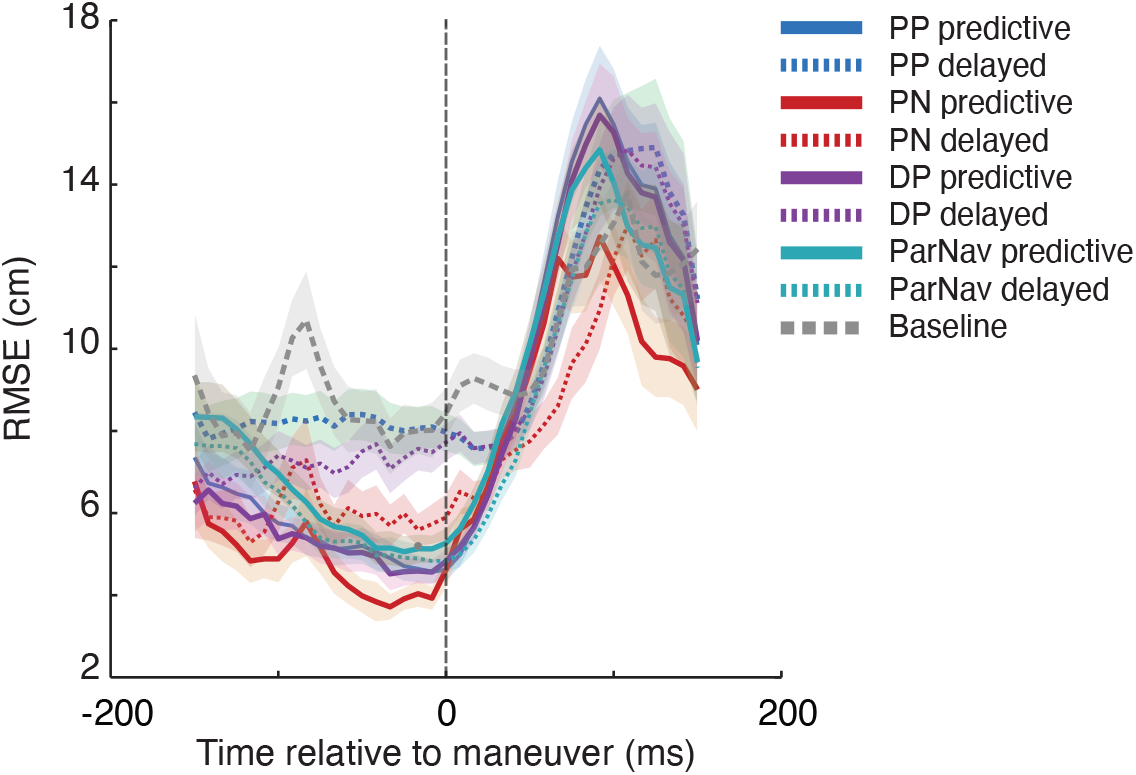
Sliding-window fit error for all eight guidance laws. Each law was fitted to the predator trajectories in sliding windows aligned to maneuver onset (dashed vertical line), with a single gain per track held fixed while every window was scored by the RMSE between simulated and observed predator positions. Line style marks the information variant (solid, predictive; dotted, delayed) and color marks the family (blue, pure pursuit; red, proportional navigation; purple, deviated pursuit; teal, parallel navigation; grey, the non-responsive baseline of a predator continuing on its committed course). Shading is the standard error across tracks; RMSE was averaged within each track before being averaged across tracks, and bins containing fewer than 15 windows were left empty. The proportional-navigation pair shown in Fig. 2b, c is the red pair here.

**Fig. 8: Extended Data Fig. 8.**
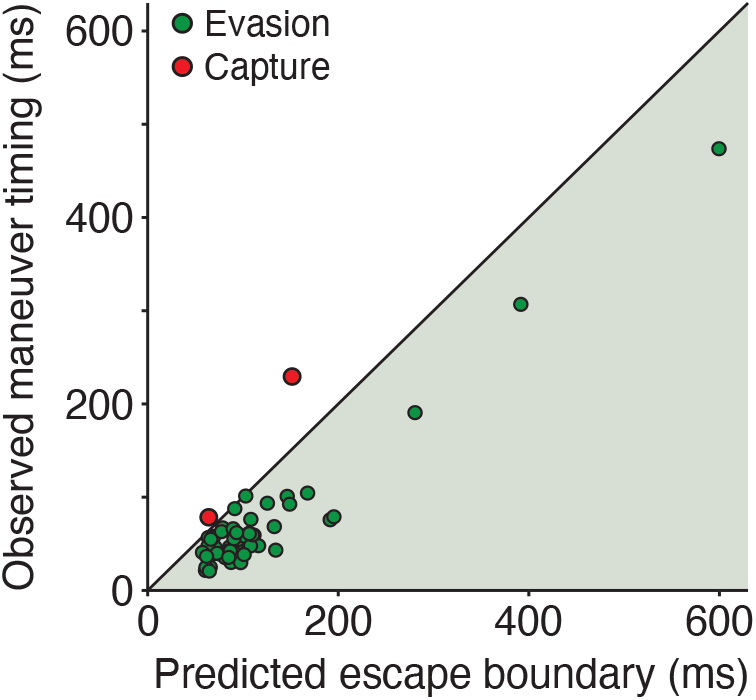
Predicted escape window for individual maneuvers. Each point is one maneuver. The predicted escape boundary is the latest time-to-collision at which the delay-extended turning gambit still allows escape, computed from the prey and predator speeds measured at that maneuver’s onset, with turning radii and sensory–motor delay at their median values. The observed maneuver timing is the time-to-collision at which the prey turned. The shaded region below the line of equality marks maneuvers initiated within their predicted escape window. Both captures (red) lie outside their window and evasions (green) inside it (*n* = 64).

**Table 1: Extended Data Table 1.**
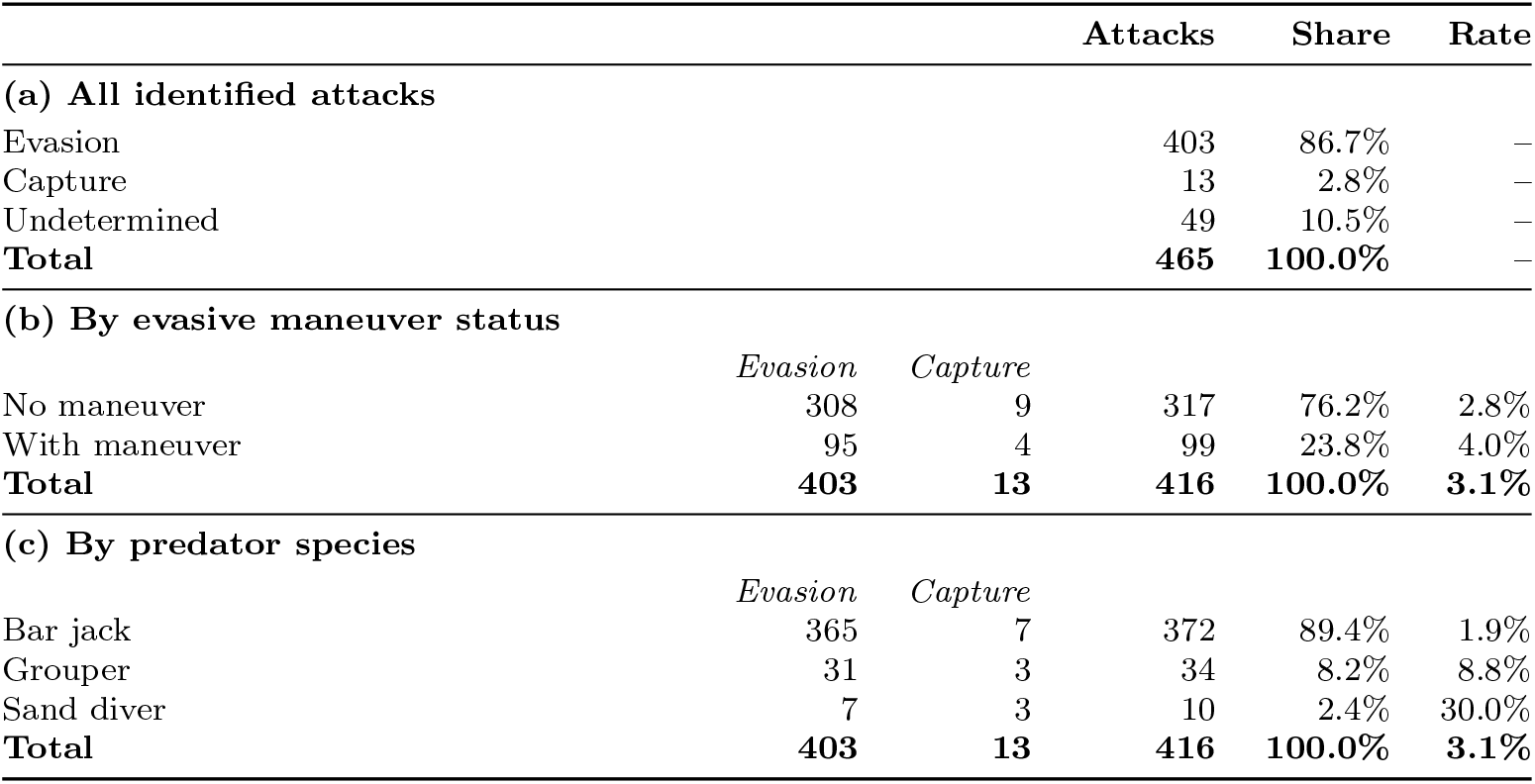
Attack outcomes and evasive maneuvers. Capture includes contact events that did not result in the prey being consumed. Panels (b) and (c) are restricted to the 416 attacks with a known outcome.

**Table 2: Extended Data Table 2.**
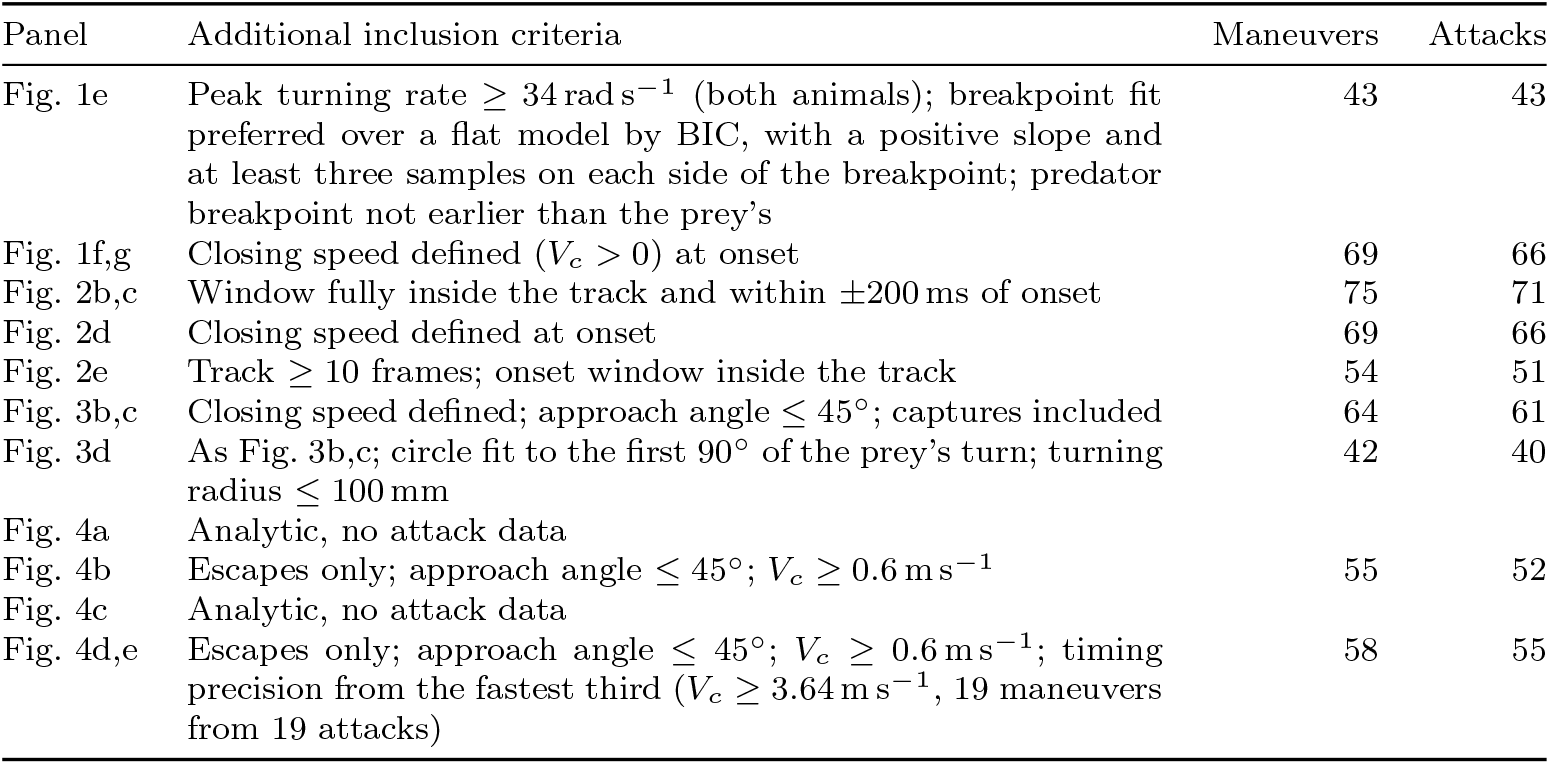
Inclusion criteria and sample size per analysis. All rows additionally require RMS 5 ≤ px (Section Field site and deployments) and start from automatically detected maneuver onsets. Fig. 2D and Fig. 3B,C each include two additional points representing captures with no detected maneuver (closing speed at the frame of contact, plotted at zero distance or TTC). Attack counts are the distinct interactions the maneuvers came from and are lower than the maneuver counts because an attack can contain more than one detected maneuver.

| Panel | Additional inclusion criteria | Maneuvers | Attacks |
| --- | --- | --- | --- |
| Fig. 1e | Peak turning rate $\geq 34 \text{ rad s}^{-1}$ (both animals); breakpoint fit preferred over a flat model by BIC, with a positive slope and at least three samples on each side of the breakpoint; predator breakpoint not earlier than the prey's | 43 | 43 |
| Fig. 1f,g | Closing speed defined ($V_c > 0$) at onset | 69 | 66 |
| Fig. 2b,c | Window fully inside the track and within $\pm 200$ ms of onset | 75 | 71 |
| Fig. 2d | Closing speed defined at onset | 69 | 66 |
| Fig. 2e | Track $\geq 10$ frames; onset window inside the track | 54 | 51 |
| Fig. 3b,c | Closing speed defined; approach angle $\leq 45^\circ$; captures included | 64 | 61 |
| Fig. 3d | As Fig. 3b,c; circle fit to the first $90^\circ$ of the prey's turn; turning radius $\leq 100$ mm | 42 | 40 |
| Fig. 4a | Analytic, no attack data |  |  |
| Fig. 4b | Escapes only; approach angle $\leq 45^\circ$; $V_c \geq 0.6 \text{ m s}^{-1}$ | 55 | 52 |
| Fig. 4c | Analytic, no attack data |  |  |
| Fig. 4d,e | Escapes only; approach angle $\leq 45^\circ$; $V_c \geq 0.6 \text{ m s}^{-1}$; timing precision from the fastest third ($V_c \geq 3.64 \text{ m s}^{-1}$, 19 maneuvers from 19 attacks) | 58 | 55 |

## Supplementary Information

### Supplementary Methods

#### Cross-eyed camera configuration

**Supplementary Figure 1:**
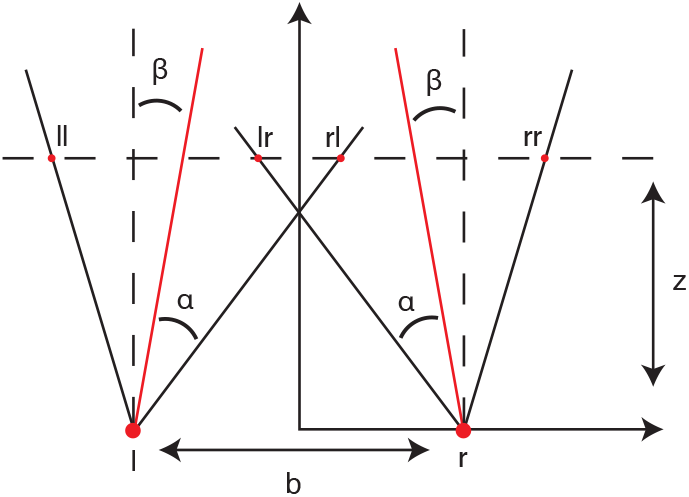
Geometry of the cross-eyed stereo camera configuration. The left (*l*) and right (*r*) cameras are separated by a baseline distance *b* and tilted inward by an angle *β* relative to the parallel-axis configuration. Each camera has a half field of view *α*. Black lines indicate the outer edges of each camera’s field of view; red lines indicate the inner edges, which cross at an intermediate distance. At distance *z* from the cameras, the four field-of-view boundaries project onto the horizontal positions *x*_ll_, *x*_lr_, *x*_rl_, and *x*_rr_ (dashed line), and the binocular overlap is the interval [*x*_rl_, *x*_lr_]. The inward tilt increases this overlap at the working distances of interest (2–5 m) compared to a parallel configuration (*β* = 0).

To maximize the overlap between the two cameras and thereby maximize the reconstructable and usable footage, the cameras were configured in a cross-eyed stereo arrangement (Supplementary Figure 1).

The two cameras of each stereo pair were mounted on a shared aluminum bar separated by a baseline distance *b*. Each camera has a horizontal field of view 2*α* (GoPro Hero 9: 2*α* = 81.3°). When the optical axes are parallel, the two fields of view begin to overlap only at some distance from the cameras, and the fractional overlap then increases slowly with *z*, remaining low at the working distances of a wide baseline. Tilting the cameras inward by an angle *β* brings the overlap closer to the cameras and increases it at intermediate distances, at the cost of reducing it at long range, where the two fields of view diverge.

For a camera pair with baseline *b*, half field of view *α*, and inward tilt *β*, the edges of each camera’s field of view projected onto a horizontal plane at distance *z* are

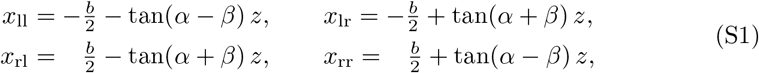

where *x*_ll_ and *x*_lr_ denote the left and right edges of the left camera’s field of view, and *x*_rl_ and *x*_rr_ those of the right camera. The fractional overlap at distance *z* is the width of the intersection of the two intervals [*x*_ll_, *x*_lr_] and [*x*_rl_, *x*_rr_] divided by the width of their union.

For *β >* 0, the overlap peaks at 100% at an intermediate distance and then declines as the two fields of view begin to diverge. The optimal tilt angle therefore depends on the working distance range. Because predator–prey interactions were observed predominantly between 2 and 5 m from the cameras, the tilt angle was chosen to maximize the mean fractional overlap over this range. Evaluating the overlap numerically for *b* = 1 m and 2*α* = 81.3° yields an optimal tilt of approximately *β* = 5°, which achieves a mean binocular overlap of 92% over the 2–5 m range, with a minimum of 80% at the near end (*z* = 2 m).

#### Three-dimensional reconstruction of the reef environment

Each interaction was placed in its local reef environment by reconstructing a three-dimensional point cloud of the reef for every attack. This serves two purposes: it provides the physical context in which the pursuit took place, and it defines the seafloor normal, which fixes the reference direction used to express trajectories in a consistent frame and to initialize the predator-centered frame (see Construction of the predator-centered coordinate frame).

A single frame from each stereo video was rectified, and landmarks were detected and matched with MATLAB’s `detectSURFFeatures` and `matchFeatures` [62]. Matches were accepted only where corresponding points fell on the same horizontal image row in the rectified views (epipolar band ≤5 px), a constraint that follows directly from rectified stereo geometry, in which true correspondences lie on identical scanlines. Accepted matches were triangulated against the stereo calibration, yielding 200–1,000 metrically scaled reference points per attack.

These points would suffice to fit the seafloor plane but are far too sparse for a scene reconstruction or for color-based segmentation, so a dense per-pixel depth map was obtained by passing the rectified left-camera image through Depth Anything V2 [63], a transformer-based monocular network that predicts depth for every pixel from a single image (Supplementary Figure 2a). Its predictions are not metrically accurate, so a linear model *Z*_metric_ = *a D*_AI_ + *b* was fitted by least squares to the sparse stereo reference points, rescaling the map to meters (Supplementary Figure 2b). The stereo points thus supply the metric scale and the Depth Anything V2 model the density. Fit quality was high and comparable in the two views: the median *R*^2^ was 0.93 for the left camera and 0.92 for the right, across all videos (Supplementary Figure 2c).

Approximately 100,000 pixels were sampled at random from the calibrated depth map and back-projected using the rectified camera intrinsics, giving a dense point cloud of the reef (Supplementary Figure 2d). The cloud was leveled by principal component analysis, taking the direction of least variance as the surface normal and rotating it onto the vertical axis, and the lowest point of the cloud defined *Z* = 0. Predator and prey trajectories were carried through the same rotation and offset, so that all reported coordinates are expressed in a frame whose vertical axis is the reef-surface normal.

For display, the leveled cloud was separated into seafloor and coral by *k*-means clustering (*k* = 2) on each point’s height and its color in CIE-Lab space. Lab was used rather than RGB because it is approximately perceptually uniform, so Euclidean distance between colors is meaningful. Features were standardized to zero mean and unit variance, height was weighted 1.2 times the color channels. The cluster with the lower mean height was taken as the seafloor (Supplementary Figure 2d, bottom).

#### Construction of the predator-centered coordinate frame

To describe the prey’s position relative to the predator, we defined a local coordinate system (***t***_*i*_, ***n***_*i*_, ***b***_*i*_) for the predator at every time step. It consists of a forward axis ***t***_*i*_ along the direction of travel, a normal axis *n*_*i*_ (up/down), and a binormal axis ***b***_*i*_ (sideways). The two transverse axes are propagated by the double-reflection rotation-minimizing frame (RMF) construction of Wang et al. [64], which introduces no rotation about the tangent beyond what the path itself forces.

The forward axis ***t***_*i*_ was estimated at each step from the change in position between neighboring samples. The normal axis was initialized along the seafloor normal (see Three-dimensional reconstruction of the reef environment), projected onto the plane perpendicular to *t*_1_, with the binormal following as ***b***_1_ = ***t***_1_ × ***n***_1_. The frame was then carried forward step by step by double reflection.

Pure parallel transport keeps the frame from twisting, but gives it no memory of any external reference. As the track curves in three dimensions the transported normal can rotate away from the seafloor normal, so that ***n***_*i*_ gradually stops meaning “up”. Each transported normal was therefore blended toward the reference direction projected into the plane perpendicular to the tangent:

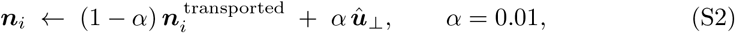

followed by re-orthonormalization against *t*_*i*_. Applied at every frame at 120 fps, the correction acts with a time constant of approximately 830 ms, long compared with the ±200 ms windows the analysis uses and short compared with the duration of a full track. Within a window the frame is therefore effectively rotation-minimizing, with under a quarter of the transported deviation pulled back. The result is a smooth frame traveling with the predator in which the prey’s bearing can be expressed as an azimuth about the predator’s heading, and in which the normal retains a consistent physical meaning throughout.

#### Guidance laws not reported in the main text

For **proportional pursuit** (**PP**), turning is proportional to the angle between the line of sight and the predator’s own heading ***û***, so(the predator g)uides steering in the direction of the prey’s position. With 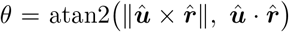, the command is a rotation about 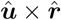,

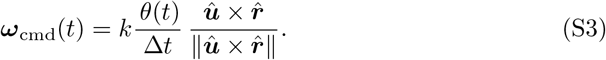

**Supplementary Figure 2:**
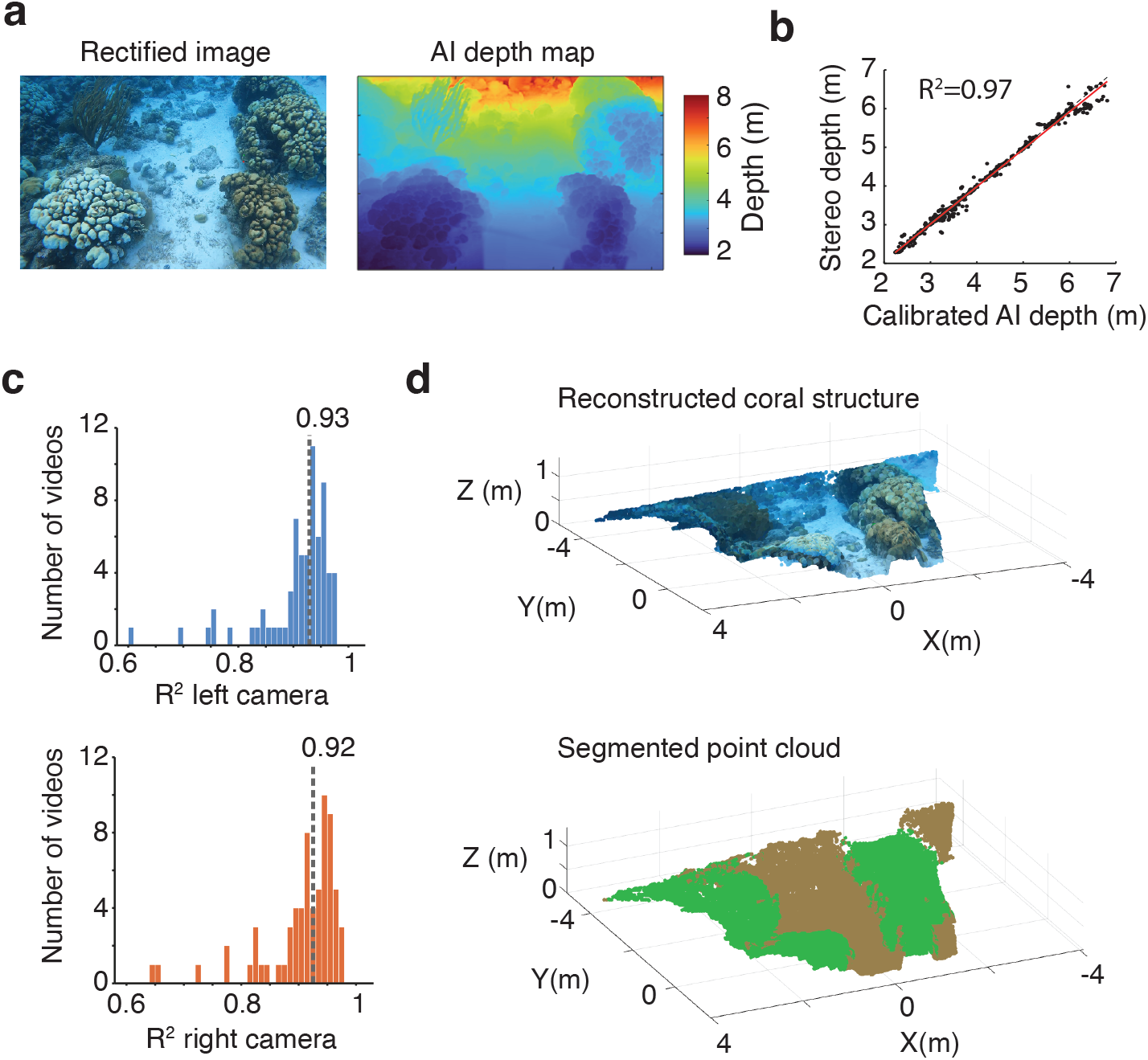
Three-dimensional reconstruction of the reef environment using calibrated AI-based depth estimation. **(a)** A rectified underwater image (left) and the corresponding metric depth map produced by Depth Anything V2 after linear calibration against stereo-derived depths (right). Color scale indicates distance from the camera in meters. **(b)** Calibrated AI depth versus stereo-derived depth for an example video, showing strong agreement (*R*^2^ = 0.97). **(c)** Distribution of *R*^2^ values for the linear depth-calibration model across all videos, for the left (blue) and right (orange) cameras. Dashed lines indicate median *R*^2^ (0.93 for left camera and 0.92 for right camera). Attacks whose prey track exceeded 5 px root-mean-square reprojection error were excluded. **(d)** Reconstructed three-dimensional coral (top) and the segmented point cloud (bottom). Green denotes coral and brown points denote sand.

Here *k* is the fraction of bearing error corrected per integration step, *k* = 1 recovers classical delay-free pure pursuit, the heading aligned with the line of sight at every step. This rule is reported for rainbow trout pursuing moving prey [21].

**Deviated pursuit (DP)** is as PP, but drives the heading–line-of-sight angle toward a fixed nonzero *θ*_*b*_ rather than to zero, so the predator settles with its heading held a constant angle off the line of sight. With *e*(*t*) = *θ*(*t*) − *θ*_*b*_ the command is

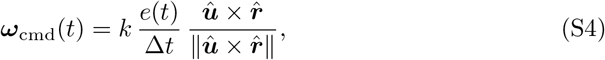

Because *θ* is unsigned, *θ*_*b*_ fixes the magnitude of the offset and not the side of the prey it falls on, so DP as implemented holds a constant angular offset rather than a directed lead. This is the strategy reported for predatory bluefish [60].

**For parallel navigation (ParNav)**, the predator holds the line of sight at a constant direction by matching the prey’s velocity component perpendicular to it and spending the remainder of its “speed budget” *s*_*p*_ closing range,

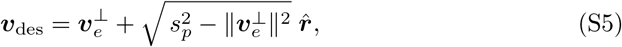

with ***v***_*e*_ as the prey’s speed vector. The desired velocity is converted to a turning rate by the rotation carrying the current heading onto it in one step,

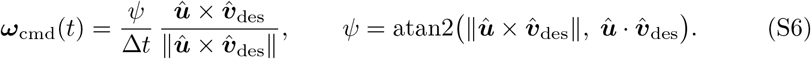

ParNav is parameter-free and was scored directly. This is the rule reported for echolocating bats [61], and the strategy against which deviated pursuit was tested in bluefish [60].

The sliding-window procedure described in Methods was applied identically to these three laws. For the pursuit laws the gain was searched over [1.2, 12]; for deviated pursuit the gain and *θ*_*b*_ were optimized jointly, with *θ*_*b*_ bounded to [0°, 90°] from a 15° start, and convergence confirmed by checking that the recovered parameters moved away from both their starting values and the search bounds. Parallel navigation is parameter-free and was scored directly.

#### Sample sizes and inclusion criteria

Of 465 identified attacks, outcome could be determined for 416 (89.5%). The remaining 49 (10.5%) were undetermined because the interaction moved out of the cameras’ field of view or behind a coral patch before it resolved. Of the 416 attacks with a known outcome, 403 (96.9%) ended in the prey evading the predator and 13 (3.1%) in capture. In three of the 13 captures the prey was partially or fully inside the predator’s mouth but was not consumed and ultimately escaped. We nonetheless classify these as captures, because the models considered here address whether the predator can bring itself into contact with a maneuvering prey, not whether it retains the prey once contact is made (Extended Data Table 1a; Supplementary Video 3).

An evasive maneuver was observed in 99 attacks (23.8% of those with a known outcome); the remaining 317 (76.2%) showed no evasive maneuver, most commonly because the prey reached coral cover before the predator came close enough to require one. Capture rate was similar between attacks with a clearly observed maneuver (4 of 99, 4.0%) and without one (9 of 317, 2.8%) (Extended Data Table 1b).

Capture rate varied by predator species (Extended Data Table 1c). Bar jack, the most frequently observed predator (372 of 416 attacks, 89.4%), had the lowest capture rate (1.9%). Sand diver, the least frequently observed predator (10 attacks, 2.4%), had the highest capture rate (30.0%). Grouper was intermediate (8.8%).

We attempted three-dimensional reconstruction for the 108 candidate attacks in which the prey performed an evasive maneuver (99) or was captured without one (9) (Extended Data Table 1b). One of these, a capture after a detected maneuver, was recorded during a short trial field season in 2022; all others are from the 2023 field season.

Reconstruction met our quality criteria for 75 of these, containing 75 automatically detected maneuvers in 71 attacks (see Evasive-maneuver detection algorithm). Reconstruction was not possible for the remainder for several reasons: stereo calibration parameters could not be determined, the interaction was visible in only one of the two cameras, or one camera drifted slightly during the deployment, invalidating its calibration for that attack.

Four of the 75 reconstructed attacks ended in capture. In two, the prey was caught after a detected maneuver. One of these is the trial capture from 2022, and in the other the maneuver carried the prey past a first predator into a second, trailing one (Extended Data Fig. 1). In the remaining two no maneuver was detected before contact, and these captures are shown at the closing speed at the moment of contact. Captures are shown separately rather than excluded wherever they appear in the figures.

Downstream analyses further restrict this pool according to the requirements of each specific test. Extended Data Table 2 gives the exact criteria and resulting sample size for every figure panel that filters attack data.

### Supplementary Note 1: Delay-extended turning gambit: derivations

#### Notation and assumptions

The delay-extended turning gambit reduces a pursuit to a small number of biomechanical quantities and a single sensory–motor delay. Its geometry and its equations of motion are introduced in the Methods (The delay-extended turning gambit). Here we derive from that model the escape boundary, the turning radius that maximizes the miss distance and the initiation distance at which the prey should begin its maneuver. The table below summarizes the notation, with subscripts *p* and *e* denoting the pursuer (predator) and evader (prey).

|  |  |
| --- | --- |
| $V_p, V_e$ | Predator and prey speeds |
| $R_p, R_e$ | Predator and prey turning radii |
| $\omega_p = V_p/R_p, \omega_e = V_e/R_e$ | Turning rates |
| $\tau_p$ | Predator sensory–motor delay, the lag between the prey’s turn and the predator’s response |
| $T_\pi = \pi R_e/V_e$ | Time the prey needs to complete its half-turn |
| $V_c = V_p - V_e$ | Closing speed before the maneuver |
| $X_0$ | Predator–prey separation at the moment the prey initiates its turn |
| $v = V_e/V_p, r = R_e/R_p$ | Speed and radius ratios |
| $\theta(t) = \min(\omega_e t, \pi)$ | Prey turn angle, from 0 at initiation to $\pi$ at the end of the half-turn |
| $\Theta(t) = \omega_p \max(t - \tau_p, 0)$ | Predator turn angle, zero until the delay has elapsed |

#### Two regimes

Which of the closed-form results derived below apply to an encounter depends on two timescales, the time the prey needs to complete its half-turn, *T*_*π*_ = *πR*_*e*_*/V*_*e*_, and the time until the predator’s own turn has carried it the prey’s final lateral offset 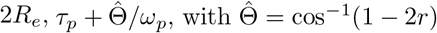 (Equation S9). When:

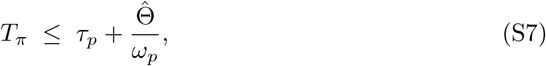

the prey has finished turning before the predator can reach its offset, so the predator can intercept it only on its straight return path, and the escape boundary has the closed form of Equation (S12). When the inequality is reversed, the predator reaches the offset while the prey is still turning, and the boundary has no closed form. The miss distances shown in the figures were computed numerically (Computation of predicted miss distance) and do not depend on this condition.

Because the escape boundary assumes the prey turns at its minimum turning radius, we evaluated this condition at that radius, 13.2 mm [10], for all the maneuvers, using its measured prey and predator speeds at onset and the median predator turning radius and delay Eq. (S7) held. The prey needed a median of 16 ms (at most 30 ms) to complete its half-turn, against a median of 46 ms before the predator could reach its offset, and the condition would fail only for prey slower than about 0.9 m s^−1^, below the slowest observed (1.4 m s^−1^).

#### The escape boundary

After completing its half-turn the prey runs straight along a path parallel to its original one but in the opposite direction, with an offset of *Y*_*e*_ = 2*R*_*e*_, one turn diameter. The predator can intercept it only once its own turn has carried it that far in the *y*-direction,

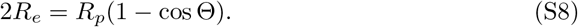

Solving it gives the turn angle at which the predator first matched the prey’s y-coordinate,

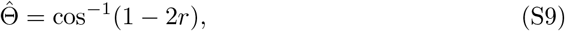

defined for 0 ≤ *r* ≤ 1, with

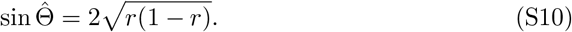

Matching *x*-coordinates (Equations 16 and 18) at the moment the predator has turned through Θ, and solving for the separation at initiation gives,

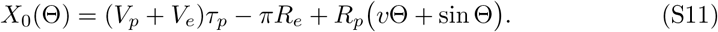

Evaluating at 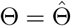 gives the escape boundary,

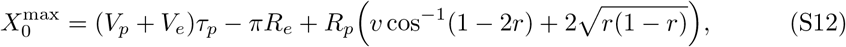

the largest initiation distance from which the prey still escapes. The corresponding time-to-collision at initiation is

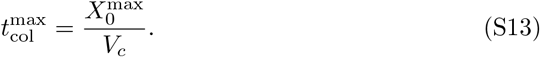

#### Miss distance at the optimal initiation distance

The prey times its escape by choosing *X*_0_, the separation at which it starts to turn. We first show that when prey maneuver at the *X*_0_ that maximizes miss distance, predator and prey reach the same x-coordinate at the point of closest approach, so that their separation in y alone sets the miss distance. Let

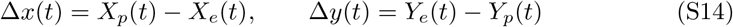

for the separations in *x* and *y*. From Equations (16)–(19), *X*_0_ enters only through the prey’s *x*-coordinate, as a constant shift, so

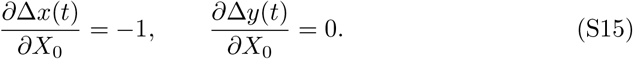

We consider initiation distances 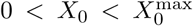, for which the prey evades the predator, so that the encounter has a well-defined point of closest approach. The positions in Equations (16)–(19) and their velocities are continuous in time, including where the prey ends its turn and where the predator starts its own, so the predator– prey distance

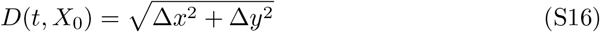

is a smooth function of both *t* and *X*_0_. The miss distance is the smallest value of *D* during the encounter, reached at time *t*^∗^(*X*_0_),

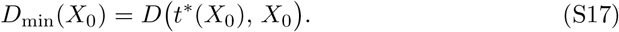

Changing *X*_0_ affects *D*_min_ both directly, through the positions, and indirectly, through the time of closest approach. By the chain rule,

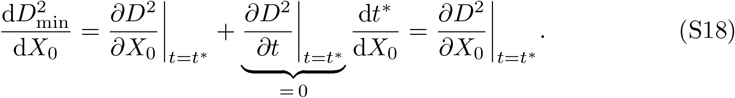

The predator is closing on the prey when the prey starts its turn, and the distance grows again once the predator’s *x*-coordinate exceeds the prey’s. So *D*^2^ first falls and then rises, and the second term vanishes at the interior minimum *t*^∗^. The first term follows from the partial derivatives above,

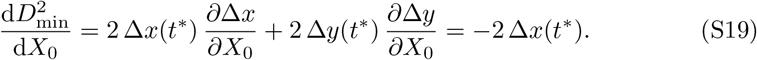

The optimal initiation distance therefore satisfies

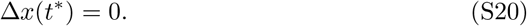

Predator and prey therefore have the same *x*-coordinate at closest approach, and the miss distance reduces to their separation in *y*,

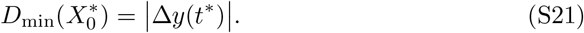

At the optimum the miss distance is purely lateral, so maximizing it reduces to maximizing the *y*-separation, which the following sections derive.

#### The *y*-separation over time

During the predator’s delay it runs straight, Θ = 0, so Δ*y* equals the prey’s own *y*-coordinate, which does not decrease. The *y*-separation can therefore only start to decrease once the predator turns, and its maximum lies at or after the end of the delay, *t* ≥ *τ*_*p*_, where

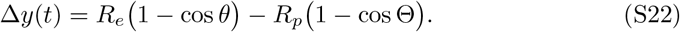

Its rate of change is the difference between the two lateral velocities,

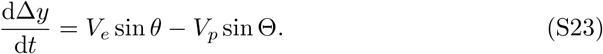

At *t* = *τ*_*p*_ this rate is non-negative, since the predator has not yet turned. By *t* = *T*_*π*_ the prey’s turn is complete and sin *θ* = 0, so the rate is strictly negative. Provided the prey is still turning when the delay ends (*T*_*π*_ *> τ*_*p*_, which holds at the optimal radius derived below, since *T*_*π*_ = *πt*^⋆^*/θ*^⋆^ *> t*^⋆^ *> τ*_*p*_), the separation therefore reaches an interior maximum on (*τ*_*p*_, *T*_*π*_), at the moment the two lateral velocities balance,

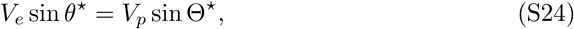

with *θ*^⋆^ and Θ^⋆^ the turn angles of prey and predator at that moment.

#### The optimal turning radius

The prey cannot turn tighter than its minimum turning radius, but it can turn wider, and doing so can increase the miss distance. At the optimal initiation distance the miss distance is the largest *y*-separation, Δ*y*(*t*^⋆^), so we ask which radius above the prey’s minimum maximizes it. By the same argument as for *X*_0_, the shift in *t*^⋆^ contributes nothing because Δ*y* is at a maximum there, so only the explicit dependence on *R*_*e*_ remains. It enters through the prey’s term in Equation (S22), via *θ* = *V*_*e*_*t/R*_*e*_, so

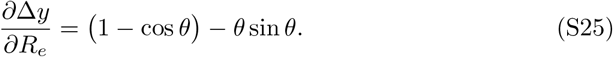

The first term is the extra room a wider turn provides, and the second is the angle the prey gives up by taking longer to complete its turn. The derivative vanishes at

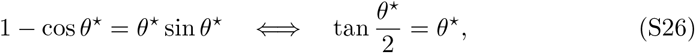

which has a single solution on 0 *< θ*^⋆^ *< π*, found numerically, *θ*^⋆^ = 2.331 rad (134°). The prey should be 134° into its half-turn when predator and prey share an *x*-coordinate.

Equation (S26) fixes the prey’s angle, and Equation (S24) then gives the predator’s,

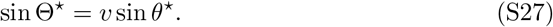

The predator reaches it at

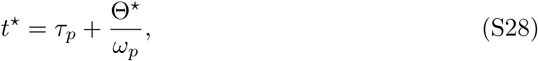

and the prey turns through *θ*^⋆^ in the same time, 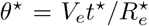. The optimal radius and the resulting miss distance are therefore

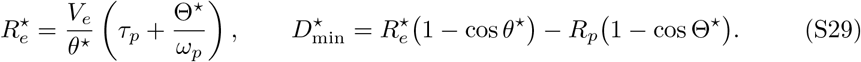

#### The optimal initiation distance

Equation (S20) requires predator and prey to share an *x*-coordinate at *t*^⋆^, which fixes *X*_0_. By then the predator has advanced *V*_*p*_*τ*_*p*_ during its delay and a further *R*_*p*_ sin Θ^⋆^ during its turn, while the prey has advanced 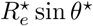 along its arc. Equating the two and eliminating 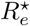 with Equations (S24) and (S29),

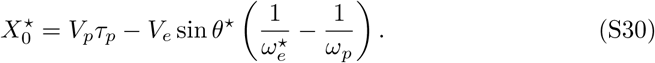

The correction is the difference in the *x*-distance the two cover while turning. It is negative when the prey turns more slowly than the predator, 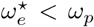, so the prey should let the predator come closer, and positive when it turns faster. It vanishes when *ω*_*p*_*τ*_*p*_ = *θ*^⋆^ − Θ^⋆^, for a predator whose turning within one delay matches the turning the prey has left to do. For the parameters considered here it is small against the first term, so

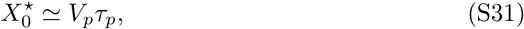

and the prey should turn approximately when the predator is one delay’s travel away, independent of the turning radii.

Writing *V*_*p*_ = *V*_*c*_ + *V*_*e*_, this decomposes as 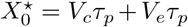, one contribution from relative closure during the delay and a second from the prey’s own displacement over the same interval. The corresponding optimal time to collision is

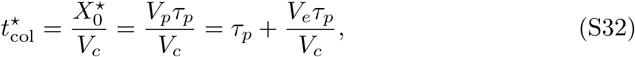

which converges on *τ*_*p*_ as the predator’s speed advantage grows.

### Supplementary Note 2: Capture probability under imprecise maneuver timing

The derivations in Supplementary Note 1 assume prey initiate at exactly the optimal moment. Here we relax that and obtain in closed form the probability that timing error carries the prey outside the escape window.

#### The escape window and its two margins

Let the prey intend to initiate at the optimal time-to-collision 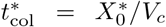 (Supplementary Note 1) but execute with a random timing error,

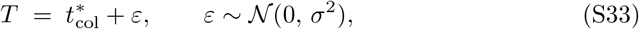

where *T* is the realized time-to-collision at onset. The miss distance is positive if and only if *T* lies in

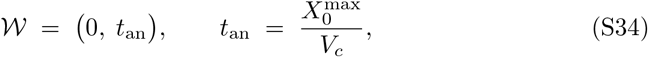

with 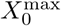 from Equation (S12). With *T* ≥ *t*_an_ the predator perceives the maneuver with enough time left to bring its course around, and with *T* ≤ 0 the maneuver does not occur before contact. Both 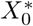 and 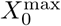 are evaluated at the prey’s optimal turning radius 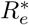 (Equation S29), which grows with prey speed. The tolerances are the distances from the optimum to each edge,

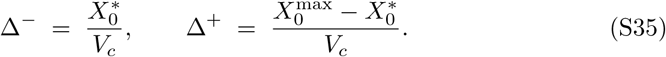

Multiplied by *V*_*c*_ both margins are distances, but they scale differently with closing speed. As *V*_*c*_ grows at fixed *V*_*e*_, *v* → 0, the predator’s turn at the optimum vanishes, sin Θ^∗^ = *v* sin *θ*^∗^ → 0, and the prey’s optimal radius approaches 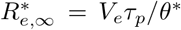. Equation (S30) then gives 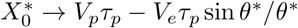, so that

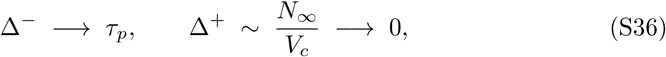

with

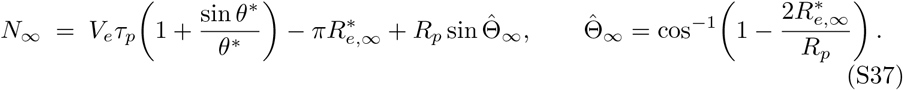

The late margin has a floor at the sensory–motor delay, whereas the early margin is a fixed distance compressed into an ever shorter time, so the window closes almost entirely from the early side.

#### Capture probability

With 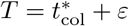, each way of failing is a condition on the timing error alone,

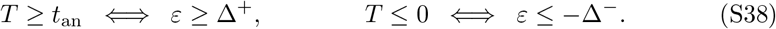

These are opposite tails of the same Gaussian and cannot occur together, so their probabilities add. Expressing each margin in units of *σ* gives

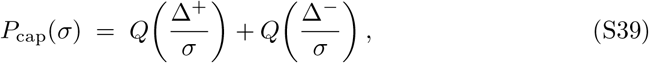

where 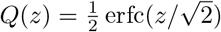 is the probability that a standard normal variable exceeds *z*. Capture probability therefore depends only on the two ratios Δ^±^*/σ*, and it increases monotonically with *σ*, from zero for perfectly precise timing toward one. Because the early margin shrinks as 1*/V*_*c*_ (Equation S36), the same timing variability carries a rising risk of capture at higher closing speeds (Fig. 4c).

### Supplementary Note 3: Evasive maneuvers are well approximated as planar

Because the delay-extended turning gambit is formulated in two dimensions, we tested whether the observed pursuit geometry justifies this approximation. For each attack, we extracted the predator and prey trajectory segments around its first maneuver, from 5 frames (42 ms) before to 10 frames (83 ms) after onset. The two segments were concatenated, centered on their shared mean and subjected to principal component analysis, so that a single plane had to describe the motion of both animals.

Across 61 maneuvers, the first two principal components captured a median of 98.6% of the positional variance and the third a median of 1.4% (IQR 0.8–2.5%; Supplementary Figure 3a, b). Because a variance fraction can be small simply because a maneuver covers a large distance, we also expressed the out-of-plane spread in absolute terms. Its standard deviation had a median of 16 mm (IQR 14–25 mm), against an in-plane extent of 141 mm, and is comparable to the 1 cm positional precision of the reconstruction (Supplementary Figure 3c). Most predator–prey interactions during the evasive maneuver are therefore approximately planar, supporting the two-dimensional approximation used in the turning-gambit derivation and the escape-boundary analysis in the main text.

**Supplementary Figure 3:**
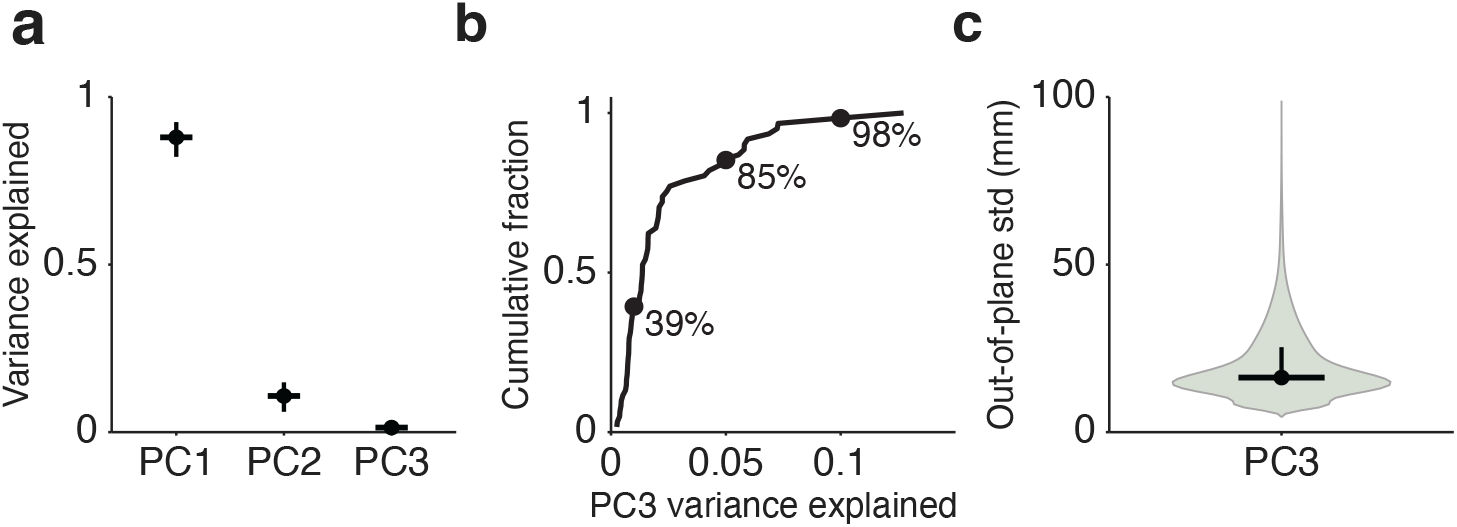
Planarity of evasive maneuvers. For each attack, the predator and prey trajectory segments from 42 ms before to 83 ms after the first maneuver were combined and analyzed by principal component analysis (*n* = 61). **(a)** Variance explained by each principal component. Dots and horizontal bars show the medians and vertical bars the interquartile range across maneuvers. **(b)** Cumulative distribution of the variance explained by the third, out-of-plane component. Markers give the share of maneuvers with less than 1, 5 and 10% of their variance out of plane. **(c)** Out-of-plane standard deviation across maneuvers (violin), with median 16 mm (dot and horizontal bar) and interquartile range 14–25 mm (vertical bar), comparable to the 1 cm positional precision of the reconstruction.

### Supplementary Note 4: Predictive control produces a turning-gambit-like maneuver

The delay-extended turning gambit assumes that, once its delay has passed, the predator turns at its maximum rate. A predator steering by a guidance law need not do so, and the escape window derived from the gambit applies to such a predator only if its controller commands that same turn. We therefore simulated a predator steering by proportional navigation, the guidance law that best described the observed trajectories (Predator guidance-law fitting), in both its reactive and predictive form with a navigation gain of *N* = 1.5, close to the median gains fitted to the predator tracks. The prey maneuvered at the optimal time for each closing speed, and we compared the resulting predator trajectories with that of the gambit (Supplementary Figure 4).

The two controllers differ in how the prey’s maneuver reaches them. The reactive controller steers on the prey’s delayed position, which changes smoothly as the prey turns, so it commands a gradual turn that stays below the predator’s limit. The predictive controller extrapolates the prey’s position from its velocity one delay earlier. When the prey’s change of heading enters that velocity, the extrapolated position shifts abruptly, the commanded turn exceeds the predator’s maximum turning rate, and the controller saturates. From that moment the predator traces the maximum-rate turn the gambit assumes, and the two trajectories coincide at every closing speed tested.

A predator using predictive proportional navigation, the controller that best described the observed trajectories (Fig. 2b, c), therefore already turns as sharply as its speed, turning radius and sensory–motor delay allow. The escape window of the turning gambit is the escape window against this predator, and no faster-turning controller could narrow it.

**Supplementary Figure 4:**
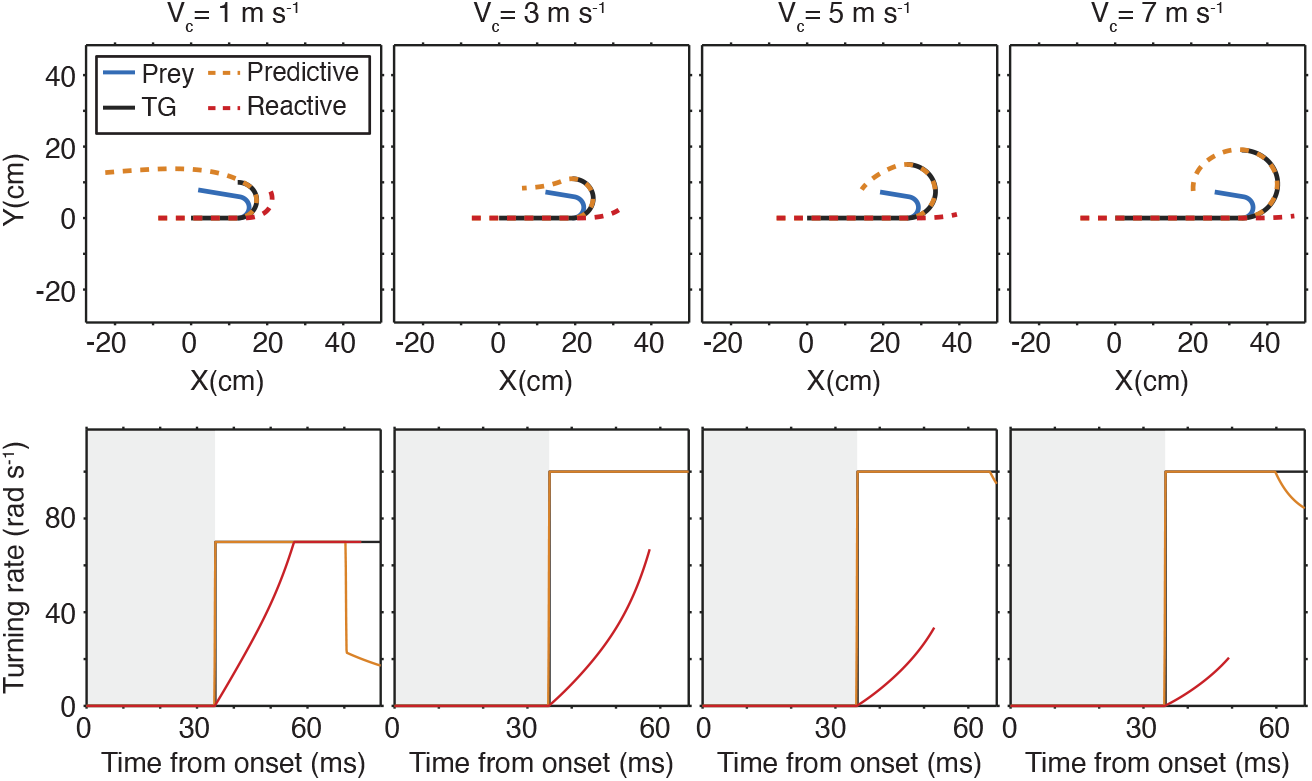
Predator trajectories under the turning gambit and under proportional navigation. Encounters simulated at four closing speeds, with biomechanical parameters at their measured medians, a navigation gain of *N* = 1.5, and the prey initiating at the optimal time for that closing speed, 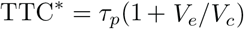. **Top**: trajectories of the prey (blue) and of the predator under the delay-extended turning gambit (black), predictive proportional navigation (orange dashed) and reactive proportional navigation (red dashed). **Bottom**: predator turning rate for the same encounters. The gray region marks the sensory–motor delay, during which the predator holds its heading. Once the delay elapses, the turn commanded by the predictive controller exceeds the predator’s maximum turning rate, so it saturates and traces the same maximum-rate turn the gambit assumes, and the two trajectories coincide. The reactive controller turns gradually instead and never reaches the limit. The ceiling differs between panels because it is the lesser of the measured maximum turn rate and the rate implied by the minimum turning radius, min(*ω*_max_, *V*_*p*_*/R*_*p*_), so the predator is radius-limited at the lowest closing speed and rate-limited above it.

